# Empirical-Bayes and Bayesian Hierarchical Modelling for Missingness and Differential Expression in Proteomics

**DOI:** 10.64898/2026.01.15.699650

**Authors:** Mengchun Li, Venkatesh Mallikarjun, Andrew Frey, Emmanuel Ogundimu, Matthias Trost

## Abstract

Mass spectrometry-based label-free proteomics data often suffer from missing values, especially for low-abundance proteins or when a protein is completely absent in one condition. This makes it challenging to estimate fold changes reliably and perform downstream analyses. Traditional imputation methods often show inconsistent performance across datasets and they typically treat imputed values as fixed rather than uncertain. This can lead to an underestimation of variability in downstream analyses. To address those problems, we present a hierarchical model that accounts for both observed protein intensities and patterns of missing data. Missing values are modelled as left-censored observations below protein-specific detection limits, reflecting the limited sensitivity of the instrument, or being missing with the probability of an intensity dependent manner. Our proposed model captures structure at multiple levels: intensity-level measurements, group-level effects (e.g., experimental conditions), and protein-level variation. To estimate model parameters, we employ an empirical Bayes framework to infer hyperparameters across proteins and use Markov Chain Monte Carlo (MCMC) methods to sample parameters from the posterior distribution. Our pipeline avoids the need for imputing missing values and is designed to produce more reliable fold-change estimates and uncertainty measures. We benchmark our method against existing approaches and demonstrate that it provides more accurate, stable, and robust estimates for differential expression analysis.

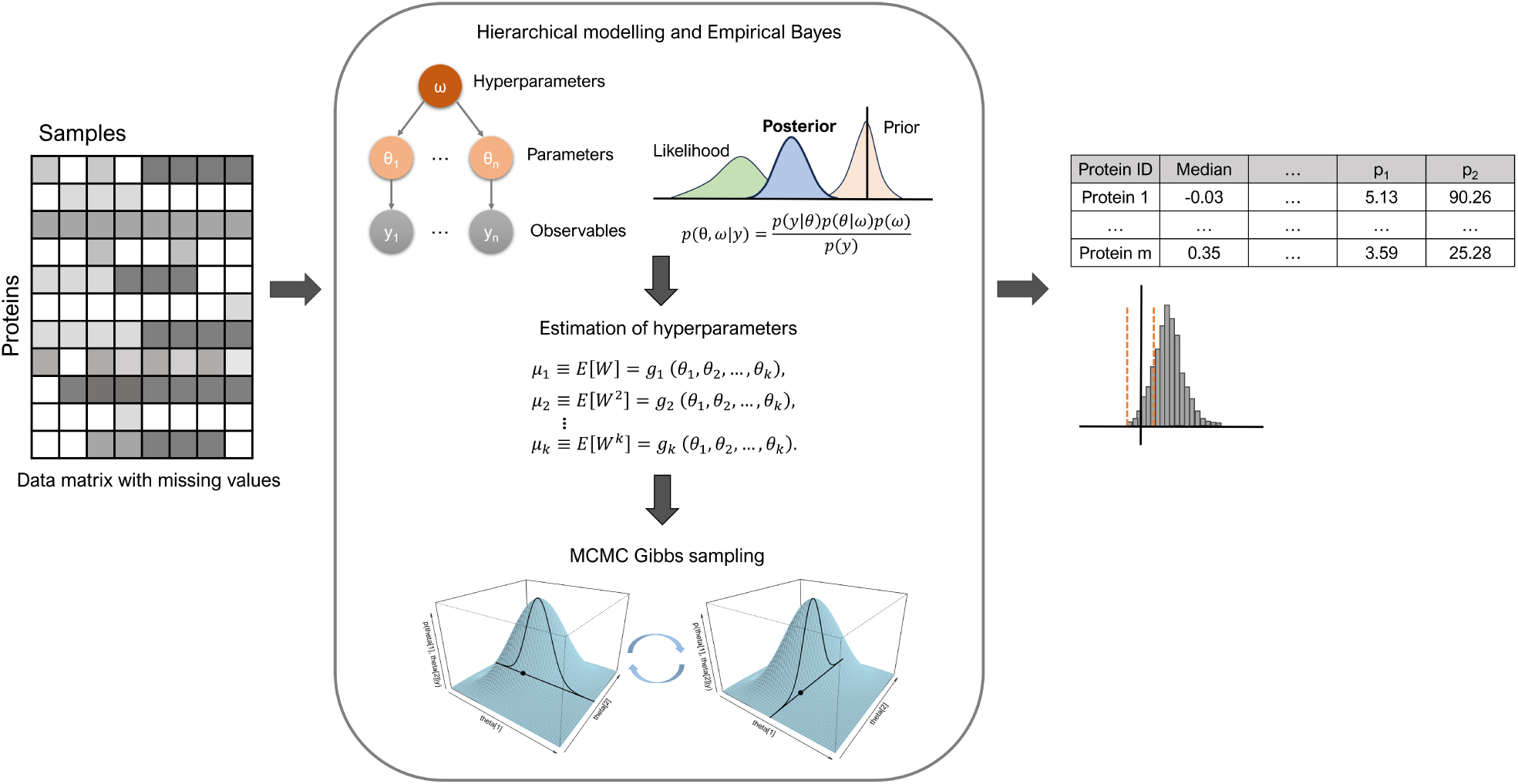

## 1 Introduction

Proteomics is the large-scale study of the proteome, including the expression, structure, functions, interactions, and modifications of proteins at any biological stage [1]. It plays a crucial role in early disease diagnosis, prognosis, and disease progression monitoring, and it is also vital in drug discovery as proteins often serve as therapeutic targets [2]. Liquid chromatography–tandem mass spectrometry (LC–MS/MS)-based proteomics enables broad detection and relative quantification of proteins [3]. Recent advances have shifted the field beyond simply identifying proteins, peptides, or post-translational modifications in small sample sets toward achieving high-quality and consistent quantification across large-scale projects involving hundreds of samples. In bottom-up proteomics, proteins are digested into peptides, separated by LC and subsequently analysed in the mass spectrometer, which provides sequence and quantitative information. Two major acquisition strategies are commonly employed: data-dependent acquisition (DDA) and data-independent acquisition (DIA) [4]. In DDA, only the most abundant (top N) ions from the MS1 scan are selected for fragmentation in the MS2 scan. Although this approach typically yields thousands of protein identifications, the stochastic selection of top N ions can introduce bias, reduce precision, and limit reproducibility by excluding low-abundance peptides. Despite these limitations, DDA remains the popular LC–MS/MS method for many proteomics studies because it is easier to implement and well-suited for relative quantification using chemical labeling techniques such as SILAC or TMT. In contrast, DIA fragments and analyzes all peptides within predefined m/z ranges during each MS2 scan, providing higher precision and reproducibility than DDA. However, the complex composite spectra generated by DIA make data interpretation more challenging [5]. Regardless of the acquisition strategy, the stochastic nature of ion selection, instrumental noise, and spectrum mismatches during library searches inevitably lead to a substantial proportion of missing values in proteomics datasets, which can pose significant challenges for downstream analysis.

Rubin [6] classified missing data into three categories: missing completely at random (MCAR), where missingness is unrelated to the data; missing at random (MAR), where it depends on other variables in an abundance-independent manner; and missing not at random (MNAR), where missingness depends on the unobserved intensity [7]. In proteomics, missing values can arise from various factors such as instrument detection limits, peptide miscleavage, low ionization efficiency, peptide co-elution, or misidentification. However, it is widely accepted that missingness is largely intensity-dependent [8–10]. This means that low-intensity peptides are more likely to be unobserved, and simply omitting these values can bias estimates upward because each missing observation still carries information about its latent intensity. Moreover, Unique proteins (those entirely absent in one condition) present an additional analytical challenge, as they allow neither fold change quantification nor conventional statistical testing. Hence, effectively addressing missing values is essential for accurate differential expression analysis in proteomics.

A common strategy to handle missing values is imputation. Various statistical methods have been proposed, such as k-nearest neighbors (kNN), random forest (RF), and singular value decomposition (SVD). However, these methods often face limitations such as the need for parameter tuning, long computation time, dependence on the missingness mechanism [11], and potential distortion of data distributions. Most imputation approaches are also deterministic, meaning they fail to propagate uncertainty into downstream inference. In typical differential proteomics workflows, imputation is followed by significance testing using t-tests, ANOVA, or moderated t-tests (e.g., `limma` [12]), all based on Null Hypothesis Significance Testing (NHST). However, such methods do not fully account for uncertainty and suffer from inherent limitations of NHST: with sufficient power, the null hypothesis can almost always be rejected, leading to inflated false discoveries [13]. Moreover, statistical significance is often misinterpreted as practical or biological relevance [14]. In contrast, Bayesian methods are more interpretable, providing direct probabilities for parameters of interest given the data and naturally incorporating uncertainty. Several EB models have been developed for differential expression analysis in proteomics, such as ProteoBayes [15] and the Selection Model for Proteomics (SMP) [9]. Despite these advances, Bayesian approaches have not yet integrated Bayesian decision-making principles or explored their practical applications in differential expression analysis. In this study, we present a hierarchical empirical Bayes model that explicitly accounts for missing values while performing differential expression analysis. We further illustrate how to interpret the posterior distribution for decision-making and evaluate model performance across different types of datasets in terms of accuracy, precision, and robustness.

## 2 Method

### 2.1 Notations

We use the following notations throughout the model:

1. *y_ijp_*: the log2 intensity of sample *i*, protein *p*, in group *j*.
2. *µ_jp_*: the mean log2 intensity of protein *p* in group *j*.
3. 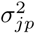: the in-group variance, which may depend on *µ_jp_* to describe sample/run effects.
4. *c_p_*: the protein-specific cutoff (shared across group), defined as the minimum observed value across all samples (*protein_min_*).
5. *µ_p_*: the overall log2 mean intensity of protein *p* across groups.
6. *σ_p_*: the between-group variance for protein *p*.
7. *µ*_1_: the mean log2 intensity across all proteins.
8. *σ*_1_: variance between proteins.
9. *α_b_, β_b_, α_p_, β_p_, µ*_0_*, γ*_0_*, γ*_1_: hyperparameters estimated from the observed data.

### 2.2 Datasets

To characterize data-distribution properties, we first analyzed two pooled datasets: a pooled liver dataset and a pooled QC dataset. Both were generated on a timsTOF platform using ddaPASEF and diaPASEF acquisition modes, and were processed with FragPipe [16] and DIA-NN [17], respectively. Complete LC–MS acquisition parameters, gradient conditions, instrument settings, and search-engine configurations are provided in the Supplementary section S1.

For model evaluation, we examined four proteomics datasets spanning both controlled and biological conditions: a controlled quantity experiment (CQE), a biological infection dataset (phagoFACS), a spike-in benchmark (UPS spike-in), and a human immune cell dataset (immune). These datasets jointly enable assessment of model accuracy, robustness, and generalizability. Detailed dataset descriptions are provided in the Supplementary table S1.

- **Pooled liver samples:** A pooled liver proteomics dataset originating from a previous study [18], acquired in DDA mode with three replicates across four conditions.
- **phagoFACS dataset:** A biological infection dataset from [19] profiling macrophage proteomes under multiple infection conditions and time points. This dataset reflects complex, real world biological variation. Data are publicly available in PRIDE under accession ID PXD062621.
- **Controlled Quantity Experiment (CQE):** The CQE dataset from [20] consists of mixtures of human, yeast, and *E. coli* proteins at known ratios, enabling direct evaluation of differential expression performance. Five conditions with five replicates in each condition were included with mixing ratios of human:yeast:*E. coli* = 35:25:40, 35:40:25, 70:20:10, 70:10:20, and 70:30:0. For benchmarking, we focused on the comparisons between 70:20:10 vs. 70:10:20 (hereafter referred to as CQE-A)and 35:25:40 vs. 35:40:25 (referred to as CQE-B), as the ground-truth fold changes are explicitly known. The dataset is available in PRIDE under accession ID PXD068192.
- **Pooled QC samples:** The pooled QC dataset was generated by repeatedly analyzing the same QC sample, which was injected 11 times across the batch using DIA. The dataset is available in PRIDE under accession ID PXD072249.
- **UPS spike-in dataset:** This benchmark dataset was originally published in [21]. It consists of a standard UPS1 protein mixture spiked into yeast cell lysate at ten concentration levels (0.01, 0.05, 0.1, 0.25, 0.5, 1, 5, 10, 25, and 50 fmol), each with four replicates. As the concentrations of the spiked proteins are known, this dataset provides a strong benchmark for evaluating precision and recall in differential expression analysis. The dataset is available in PRIDE under accession ID PXD009815.
- **Immune dataset:** From [22], this dataset profiles 28 primary human hematopoietic cell populations under steady and activated states. The dataset is available in PRIDE under accession ID PXD004352.

### 2.3 Proteomics data distribution

#### 2.3.1 Data distribution

We examined four proteomics datasets, including both DDA and DIA data. Because raw protein intensities are typically right-skewed, log-transformation is commonly applied to stabilize variance, reduce skewness, and make the distribution more symmetric and normal-like. Therefore, we used log2-transformed intensities throughout our analysis. Unless otherwise specified, all intensity values reported in this study refer to log2-transformed protein intensities. The pooled liver dataset (Figure 1A) is a DDA dataset with 21.4% missing values, while the phagoFACs, CQE and pooled QC datasets (Figure 1B-D, respectively) are DIA datasets with missing proportions of 18.2%, 11.7%, and 1.1%. The blue area represents the observed data, and the red curves correspond to normal distributions fitted to the right-hand side of the observed distributions. As the proportion of missing values decreases, the observed distributions appear increasingly normal, supporting the assumption that the underlying true distributions follow a normal pattern. Additionally, missing values are concentrated at the lower end of the distribution, indicating that missingness is intensity-dependent: proteins with lower intensities are more likely to be missing.

**Figure 1:**
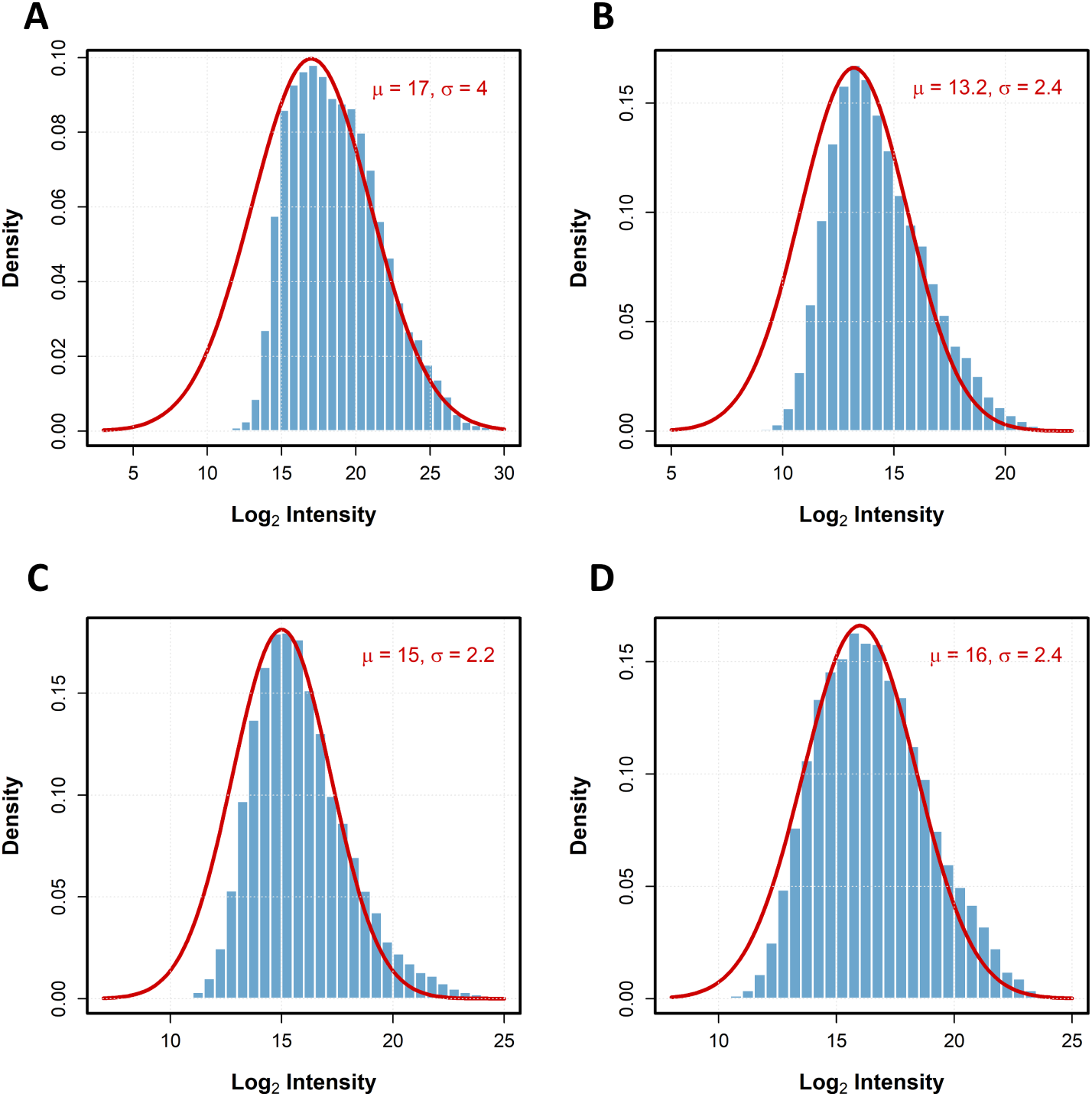
Protein intensity distributes near normally and missing values are intensity dependent. Observed protein intensity distributions for four proteomics datasets (A–D) with missing value proportions of 21.4%, 18.2%, 11.7%, and 1.1%, respectively. Blue areas represent observed data, and red curves show normal distributions fitted to the right-hand side of the observed distributions. The x-axis displays log2 intensities; densities are normalized to 1 and scaled by the proportion of observed values. (A) pooled liver dataset. (B) phagoFACS dataset. (C) CQE dataset. (D) pooled QC dataset.

#### 2.3.2 Missing pattern

As noted previously, missing values in proteomics datasets tend to be intensity-dependent. Some studies have proposed that the detection probability of a protein intensity follows a probabilistic dropout function, such that the number of detected samples per protein can be modeled by a zero-truncated binomial distribution [10,23].To investigate this, we analysed the log2 intensity of the phagoFACS dataset which has six replicates per group. For each protein, we first calculated the group-wise sample means 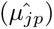, then binned these means in intervals of width 0.5. We assumed that proteins from different groups whose means fall within the same interval share the same detection probability. Next, we counted the number of observed values for the protein groups within each interval. If the in-group samples (each trial) are independent and identically distributed, the observed counts in each interval should approximately follow a zero-truncated binomial distribution. Our results (Figure 2) indicate that for proteins with higher group means (above 14), the observed counts closely follow a zero-truncated binomial distribution, with most protein groups showing no missing values. However, at the lower intensities (means below 14), we observed more protein groups with complete observations and fewer with one missing value than would be expected under a purely intensity-dependent logistic model. This indicates that missingness is not solely driven by an intensity-based probability, the detection process is conditionally independent but not identically distributed. Based on these findings, we hypothesize that missingness can also arise from protein-specific cutoffs, reflecting differences in ionization efficiency, chromatographic behaviour, or fragmentation patterns. Although mass spectrometers have a global limit of detection (LOD), in practice, each protein may have its own effective detection threshold below which its signal cannot be distinguished from background noise [9]. Proteins with relatively low cutoffs have intensities that remain above their detection threshold in all replicates and therefore appear to be fully detected. Nevertheless, even above these cutoffs, random missingness can still occur due to factors such as ion interference or peak-picking errors. To account for these two sources of missingness, we introduced a hybrid modelling strategy. Specifically, proteins with a high missing proportion (above a threshold) are modelled using a cutoff-based mechanism, while those with a low missing proportion are modelled using a logistic dropout function. The effect of varying this threshold on model performance is investigated in the Results section.

**Figure 2:**
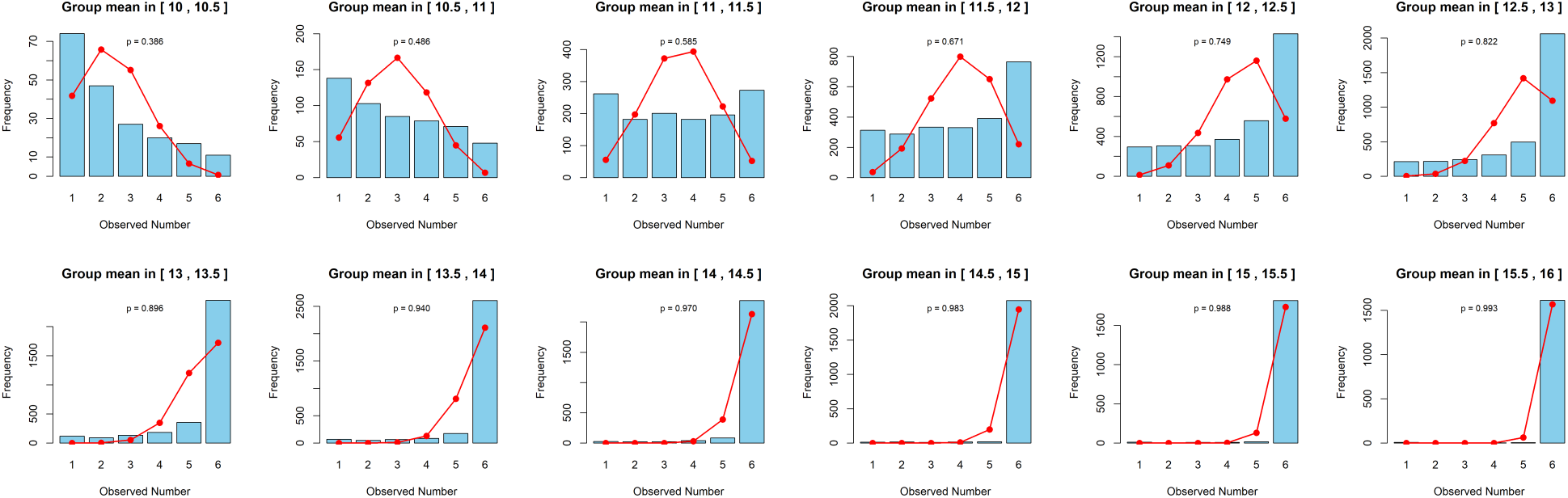
Distribution of missingness within protein groups with similar sample means. Blue bars represent the observed number of missing values per protein per group within each intensity interval, and the red curve shows a zero-truncated binomial distribution fitted using maximum likelihood estimation (MLE). Deviations at lower intensities indicate that missingness cannot be fully explained by a simple intensity-dependent logistic model.

### 2.4 Assumptions

The EB model relies on several key assumptions. First, within each group, the observed protein intensities *y_ijp_* are assumed to follow a normal distribution with mean *µ_jp_* and standard deviation *σ_jp_*. At the group level, the mean intensities *µ_.p_* are also assumed to be normally distributed, with mean *µ_p_* and standard deviation *σ_p_*. Across all samples, the true underlying protein intensities are modeled as normally distributed with mean *µ*_0_ and standard deviation *σ*_0_. Finally, missing values are assumed to occur either due to protein-specific detection cutoffs or at random with a probability that depends on the latent intensity (intensity-dependent missingness).

### 2.5 Model

#### 2.5.1 Hierarchical modeling

Let *y_ijp_* denote the latent (true) log_2_ intensity of protein *p* in group *j* and sample *i*. We assume

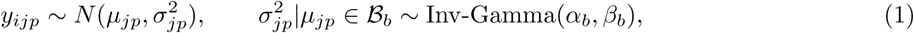

where ℬ*_b_* is the b-th mean-intensity bin.

Observed values are generated from the latent intensities through a detection mechanism. Option 1 treats intensities below a protein-specific threshold *c_p_* as left-censored. We define

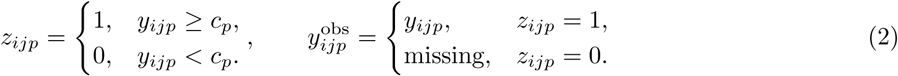

The corresponding censored-data likelihood for protein *p* in group *j* is

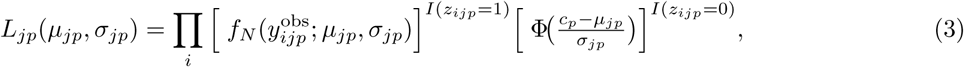

where *f_N_* (·; *µ, σ*) is the normal pdf. This formulation assumes *c_p_* is approximately constant across experimental conditions.

Option 2 models the probability of detection as a logistic function of the latent intensity,

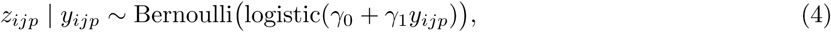

the parameters *γ*_0_ and *γ*_1_ are global coefficients. *γ*_1_ controls how detection probability increases with intensity and *γ*_0_ is the intercept of the log-odds as a function of intensity. To estimate *γ*_0_ and *γ*_1_, we fit a logistic regression model using the observed intensities and their estimated detection proportion. Because the proportion of missing values at a given intensity cannot be observed directly, we approximate the number of missing values at each intensity level by exploiting the empirical symmetry of the latent distribution: for each observed intensity bin, we identify a corresponding bin on the opposite side of the distribution mean, and use the excess frequency in that symmetric bin as an estimate of the number of missing values. The resulting estimated detection proportions are then used as the response in a logistic regression (R function `glm`).

For observed values the likelihood is the product of the normal pdf and the detection probability evaluated at the observed *y*, whereas for missing values the likelihood is the marginal probability of being undetected,

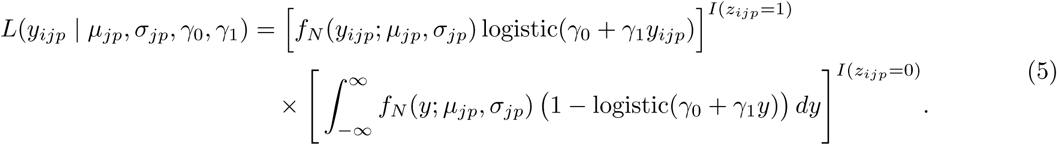

For j experimental conditions with group means *µ*_1*p*_*,…, µ_jp_*, we assume a hierarchical prior:

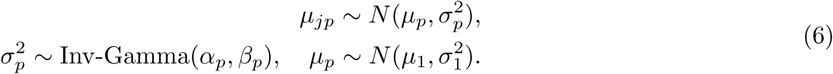

This hierarchical structure shrinks group means toward the overall protein mean, reducing false positives while leveraging the assumption that most proteins remain unchanged across conditions.

By marginalizing over groups, the expected distribution of *y_ijp_* is

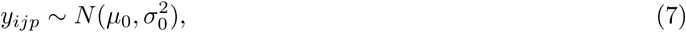

where *µ*_0_ = *E*[*y_ijp_*]. Given

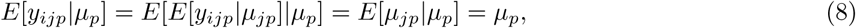

therefore

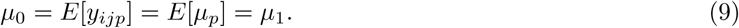

The hierarchical structure of the EB model is illustrated in Figure 3. This diagram summarizes the dependencies among observed intensities, group-level means, protein-level means, and hyperparameters, highlighting how information is shared across groups and proteins to improve estimation accuracy.

**Figure 3:**
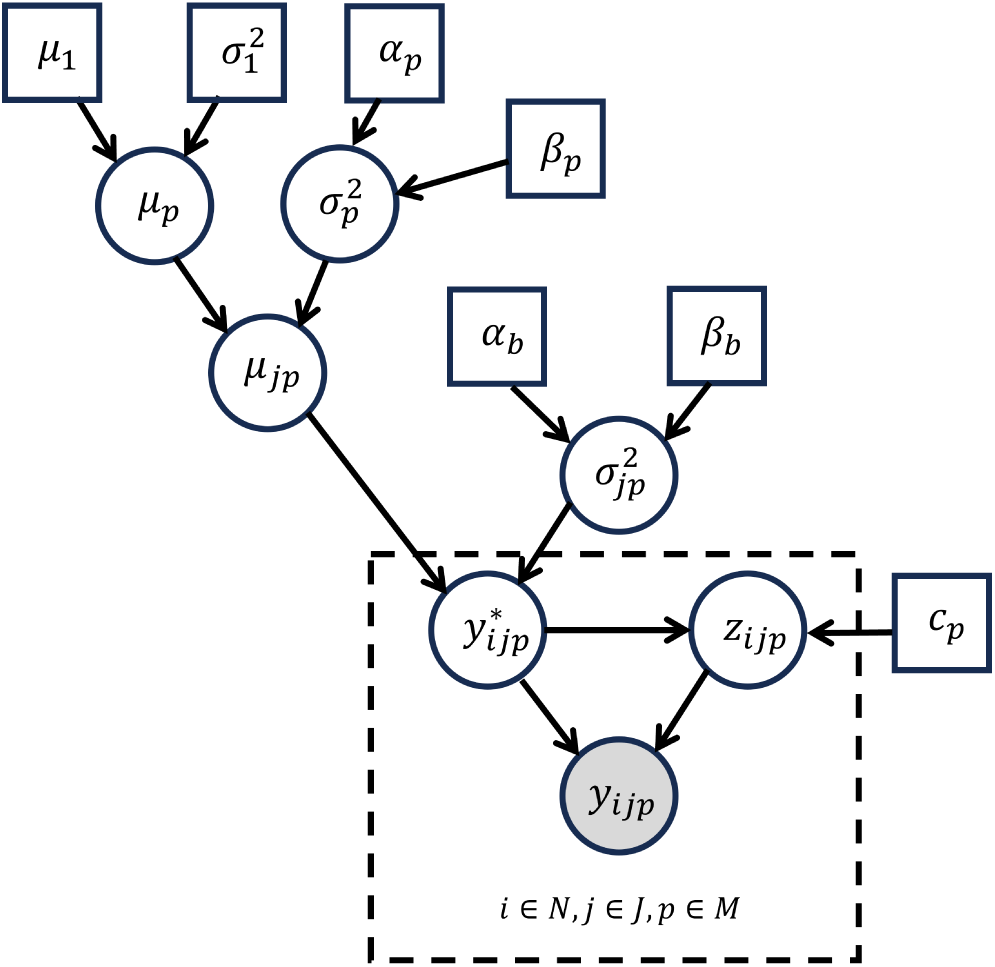
Probabilistic graphical model for the EB model under the censored-normal missingness mechanism. Nodes represents N observations across J groups and M proteins. Shaded circles correspond to observed intensities, solid circles to latent stochastic variables, solid rectangles to fixed constants, and dashed rectangles indicating replication. Notation follows [24].

#### 2.5.2 Mean-variance trend

We modelled the in-group variance using an inverse-gamma distribution. Previous studies have reported the mean–variance trend, which describes the relationship between a protein’s mean intensity and its variance across samples [25, 26]. In line with this concept, we accounted for the dependence of variability on mean intensity. However, rather than modelling variance as a smooth deterministic function of the mean, we modelled the distribution of variances conditional on the mean. In this framework, given the mean abundance of a protein group, the variance is not restricted to a fixed curve but is instead treated as a random variable drawn from an inverse-gamma distribution parameterized by the mean, which integrates naturally into our hierarchical model. Empirically (Figure 4), we observed that in-group standard deviations tend to be higher at lower mean intensities and become more compact at higher mean intensities. To capture this trend, we partitioned protein groups into bins according to their in-group mean intensity and, within each bin, estimated the parameters of an inverse-gamma distribution for the in-group variances using the method of moments. A detailed derivation is provided in the Supplementary section S2.

**Figure 4:**
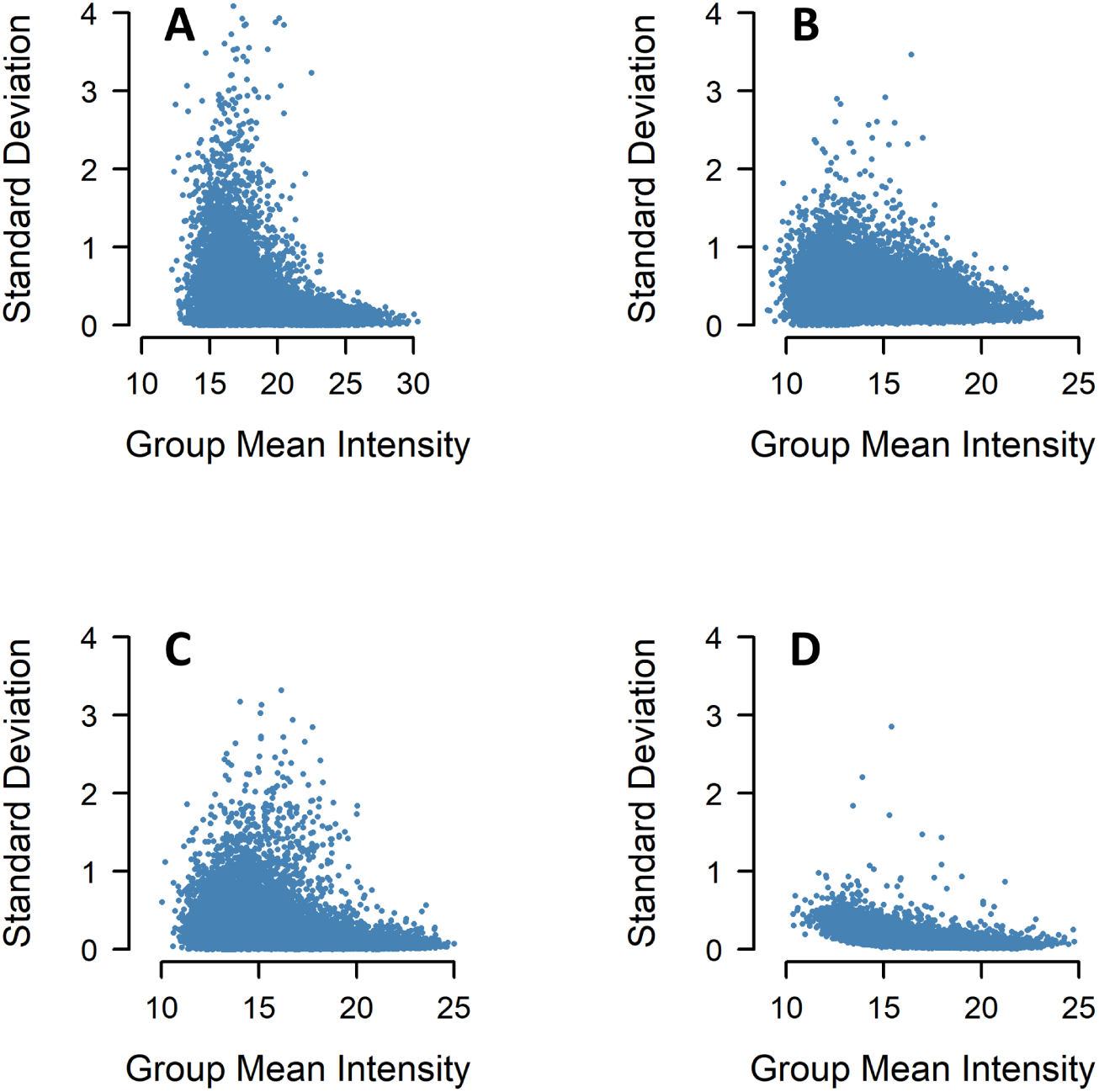
Mean-variance trend of four different datasets. Each point represents a protein within a group, with the group mean intensity plotted against the corresponding group standard deviation. Standard deviations are more dispersed at lower mean intensities and become more stable at higher intensities, indicating a mean-dependent variance structure. (A) pooled liver dataset. (B) phagoFACS dataset. (C) CQE dataset. (D) pooled QC dataset.

#### 2.5.3 Bayesian Inference

According to Bayes’ theorem, the posterior distribution of model parameters *θ* given data *D* is *p*(*θ*|*D*) = *p*(*D*|*θ*)*p*(*θ*)*/p*(*D*). In our case, the primary quantity of interest is the marginal posterior distribution of the log fold change, expressed as *p*(*µ*_1*p*_ − *µ*_2*p*_ | *D*). To obtain this distribution, we performed Bayesian inference using Markov Chain Monte Carlo (MCMC) sampling, which generates posterior distributions by iteratively drawing samples from conditional probability distributions. This approach allows us to approximate the joint posterior of model parameters without requiring closed-form solutions [27]. Model fitting was implemented in rjags, an R interface to the JAGS (Just Another Gibbs Sampler) software [28], which provides a flexible framework for specifying hierarchical EB models and running MCMC simulations. We monitored accuracy and convergence using standard diagnostics (e.g., ESS, Gelman–Rubin statistics) and based all posterior summaries on the combined draws from multiple independent chains.

#### 2.5.4 Estimation of hyperparameters

The hyperparameters were estimated using an empirical Bayes (EB) approach. Specifically, the parameters *µ*_1_ and 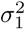 were obtained by maximizing a censoring-aware marginal likelihood of the protein-level means. The global prior mean for latent intensities was set as *µ*_1_ = *µ*_0_.

For the variance, inverse-gamma hyperparameters *α_b_* and *β_b_* were estimated across proteins within bins defined by their mean intensity, allowing the mean–variance trend to be captured in a bin-specific manner. These EB plug-in estimates were then treated as fixed hyperparameters in subsequent hierarchical modeling.

#### 2.5.5 Decision making and FDR control

We use the concept of the region of practical equivalence (ROPE), which specifies a small range of parameter values considered practically equivalent to the null value for the application of interest [29]. For example, when testing whether a protein is differentially expressed, we are primarily interested in whether the underlying log fold change (logFC) is close to zero. Small deviations above or below zero are practically negligible.

We define the hypotheses and possible actions as follow:

*H*_0_: The protein is not differentially expressed,
*H*_1_ : The protein is up-regulated,
*H*_2_ : The protein is down-regulated.
*A*_0_ : declare null,
*A*_1_ : declare up,
*A*_2_ : declare down.

From posterior distribution we calculate

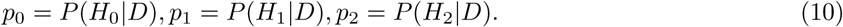

We define the loss matrix as

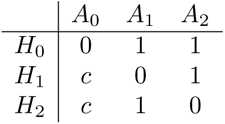

where *c* ≥ 0 represents the cost of a false negative (i.e., failing to detect a true differential expression compared to making a false positive decision).

The expected losses are therefore

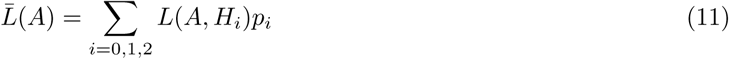

The Bayes decision rule is to select the action *A_k_* that minimizes the posterior loss:

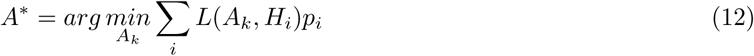

In case of ties where two actions yielding the same expected loss, we follow a conservative rule and choose the null decision *A*_0_. This formulation follows standard Bayesian decision analysis [30], ensuring that both false positives and false negatives are explicitly weighted through *c*.

Local false discovery rate (local FDR), also called the posterior error probability (PEP), is the probability that a declared positive is a false discovery given the observed data [31]. For declaring *A*_1_ (up-regulated), this is

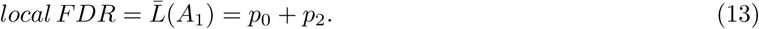

In our Bayesian decision framework, the optimal decision rule can be interpreted in two equivalent ways, depending on how the cost ratio *c* is specified:

1. Cost-sensitive Bayes rule. When minimizing the posterior expected loss, declaring *A*_1_ is optimal if

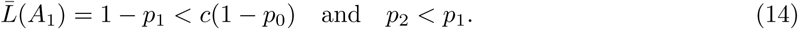

or equivalently,

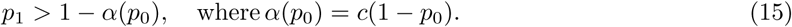

Here, the threshold *α*(*p*_0_) varies with *p*_0_.
2. Fixed local-FDR cutoff. Alternatively, one may impose a constant local FDR threshold by declaring *A*_1_ when *p*_1_ *>* 1 − *α*, where *α* is a fixed cutoff controlling the posterior probability of false discoveries.

To maintain consistency, the decision framework throughout this paper adopts the fixed local-FDR cutoff formulation (option 2), where *α* = 0.05 is chosen to control the desired posterior false discovery rate. Let *S_up_* and *S_down_* be the sets of proteins declared up- and down-regulated, with *N* = |*S_up_*| + |*S_down_*|. The Bayesian global FDR can then be defined as

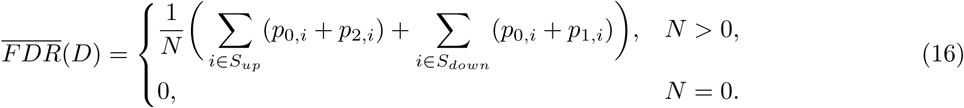

In practice, however, multiple testing requires us to control the frequentist expectation of FDR, which represents the proportion of false positives among all declared positives. Müller et al. established FDR guarantees for Bayesian procedures under mixture priors with a point mass at the null [32]. Our model uses a continuous prior on fold changes, so those formal guarantees do not strictly apply. However, the ROPE framework assigns positive prior probability to a null region | log FC| *< δ*, which is analogous in spirit. We therefore assess FDR control empirically (Figure 5), comparing posterior-based (“theoretical”) FDR with the actual FDR across ROPE widths ([−*δ, δ*], with *δ* from 0.1 to 0.5) and significance thresholds *α* = 0.1, 0.05, 0.01, using the CQE dataset comparison A. The theoretical FDR tends to underestimate the actual FDR when the ROPE is narrow, indicating a slightly optimistic model and to overestimate it when the ROPE is wide, reflecting a more stringent decision rule. This pattern is intuitive: widening the ROPE reduces the likelihood of rejecting *H*_0_, thereby decreasing the actual FDR but increasing the theoretical FDR because a larger ROPE region contributes more directly to its calculation. Although the two FDR measures are not identical, both consistently remain below their corresponding *α* thresholds, demonstrating that our model effectively controls the FDR below the desired level.

**Figure 5:**
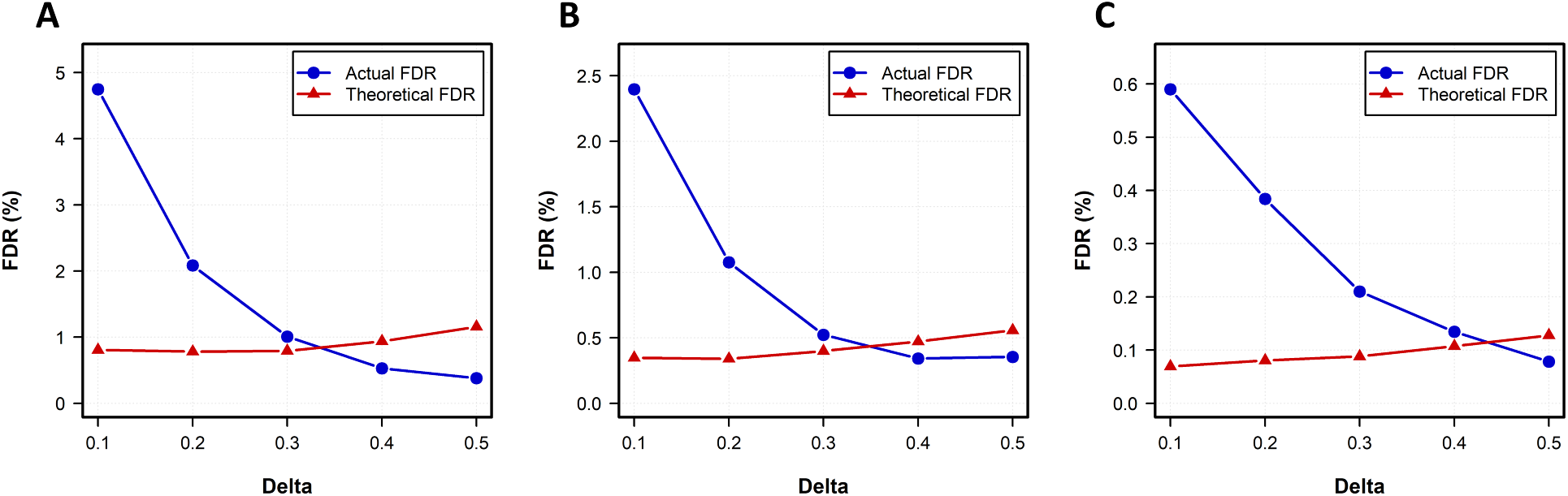
Comparison of actual and theoretical FDR across ROPE widths. (A) *α* = 0.1. (B) *α* = 0.05. (C) *α* = 0.01. The blue curves show the actual FDR, calculated as the proportion of false positives among all detected positives. The red curves show the theoretical (posterior-based) Bayesian FDR. Although the two measures exhibit different trends, they both consistently remain below their respective target *α* levels, indicating effective FDR control by the model.

Both the ROPE and the decision thresholds (*c* or *α*) determine the stringency of rejection. However, it is important to distinguish their roles. The parameter *c* is a cost ratio in the loss function: it represents the relative penalty of failing to detect a truly differentially expressed protein versus incorrectly declaring a protein as differential expressed. By contrast, *α* is a fixed local-FDR cutoff that controls a posterior error probability. In practice we therefore recommend the following workflow. First, choose a ROPE width consistent with what effect sizes are scientifically meaningful. Second, select a small set of plausible cost ratios *c* (for example *c* ∈ {0.5, 1, 2, 5, 10}) that represent trade-offs for the study (e.g. based on cost or feasibility of downstream analysis). Alternatively, evaluate a few fixed local-FDR cutoffs *α* (e.g. 0.01, 0.05, 0.1). Report numbers of discoveries and estimated 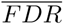 across these values and choose a final threshold to be pre-specified in protocols depending on the study goals.

#### 2.5.6 Output interpretation

To illustrate the output of our empirical Bayes (EB) model, we present results for three representative proteins from the UPS spike-in dataset. Table 1 summarizes the posterior statistics, and Figure 6 shows the posterior distributions of the log fold changes. The highest density interval (HDI Low and HDI High) marks the bounds of the posterior distribution’s highest density interval, which by default is set at 95%. The Median reflects the central tendency of the distribution. The null value (0) divides the distribution into two parts; pLtCompVal reports the smaller tail probability, indicating how far the distribution is from the null. The posterior probabilities *p*_0_, *p*_1_ and *p*_2_ defined earlier correspond directly to the proportions of the posterior distribution lying within, above and below the ROPE. Specifically, *pInROPE* = *p*_0_, *pGtROPE* = *p*_1_, *pLtROPE* = *p*_2_. These three probabilities represent exclusive regions of the posterior and therefore satisfy the sum equals to 1. For example, P00167 shows a 99.94% probability of being down-regulated, only 0.05% probability of no change, and 0.01% probability of up-regulation. In contrast, 89.67% of the posterior mass for O13563 lies within the ROPE, supporting the conclusion that it is not differentially expressed. For P03874, 97.45% of the posterior lies above the ROPE, suggesting up-regulation. It is also noteworthy that the HDI intervals for P00167 and P03874 are wider than for O13563. This is because both have one condition entirely missing, whereas O13563 has complete data, making its estimates more precise.

**Figure 6:**
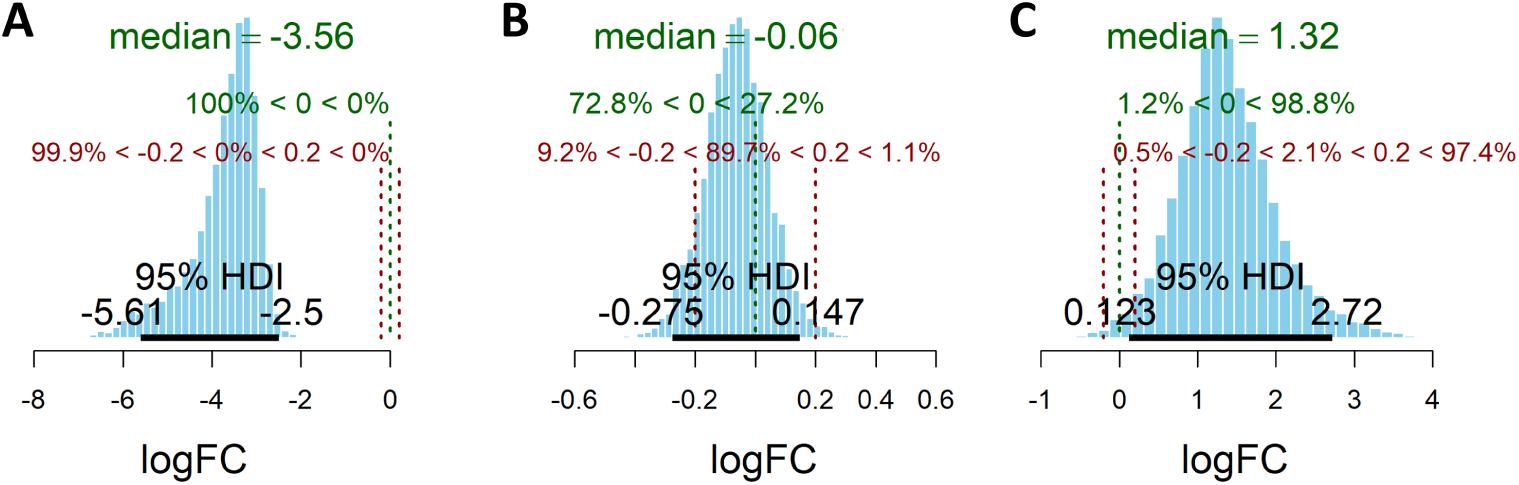
Posterior distributions of log fold changes for three representative proteins from UPS spike-in dataset: (A) P00167 (down-regulated). (B) O13563 (no differential expression). (C) P03874 (up-regulated). Shaded regions show the posterior distributions; red vertical lines denote the ROPE boundaries, and the green vertical line marks the null value (0). Percentages of posterior mass falling between key values are displayed on each plot. The 95% highest density interval (HDI) is indicated by the black line along the x-axis.

**Table 1:** Posterior summary for representative proteins.

| Protein ID | HDI.Low | HDI.High | Median | pLtCompVal(%) | pLtROPE(%) | pInROPE(%) | pGtROPE(%) |
| --- | --- | --- | --- | --- | --- | --- | --- |
| P00167 | -5.68 | -2.41 | -3.59 | 0.04 | 99.94 | 0.05 | 0.01 |
| O13563 | -0.28 | 0.15 | -0.06 | 27.17 | 9.20 | 89.67 | 1.13 |
| P03874 | 0.12 | 2.72 | 1.32 | 1.25 | 0.48 | 2.07 | 97.45 |

## 3 Validation

### 3.1 Competitive Methods

Since imputation performance can be strongly affected by factors such as sample size, data distribution, and the proportion of missing values [33], we compared our model against a diverse set of established imputation strategies. The goal was to cover the major approaches used in proteomics, including simple single-value methods, local correlation–based methods, global low-rank approaches, and specialized tools, to ensure a fair and comprehensive evaluation across different assumptions and complexity levels. In addition to these imputation strategies, we included `limma` as a benchmark differential expression (DE) analysis method. Throughout this manuscript, “limma” refers specifically to the moderated t-test applied to the observed (non-imputed) data. For all other imputation methods listed below, missing values were first imputed, and the resulting completed datasets were analyzed using `limma`.

- **Single-value imputation:** imputes all missing values with a fixed value, such as replacing missing values with the lowest observed intensity of each variable (LOD) or values drawn from a left-shifted normal distribution (ND) [34]. These methods are simple and computationally efficient but often underestimate variance.
- **Local similarity:** methods such as k-nearest neighbors (kNN), local least squares (LLS), multivariate imputation by chained equations(mice-pmm) and random forest (RF) imputation leverage the similarity among subsets of proteins or samples to predict missing values [35].
- **Global similarity approaches:** methods such as Bayesian principal component analysis (BPCA), singular value decomposition (SVD), and maximum likelihood estimation (MLE) decompose the data matrix or minimize the determinant of the covariance and reconstruct missing values accordingly [36].
- **Other specialized methods:** for example, `msImpute` [37], which has been specifically developed for mass-spectrometry proteomics data and accounts for missing not at random (MNAR) values.

### 3.2 Performance metrics

- RMSE is the average error between estimated fold change and true fold change for all proteins with missing values.

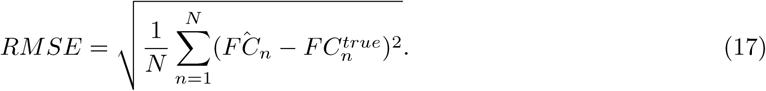

- Precision is the proportion of proteins that are predicted as TP among all predicted positives.

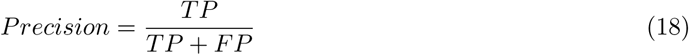

- Recall is the proportion of proteins that are predicted as TP among all actual positives.

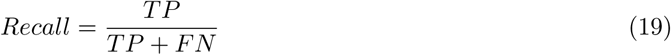

- False sign rate is defined as

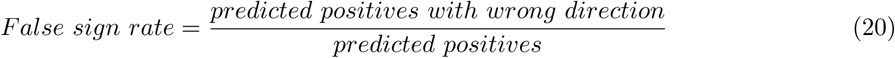

- F1 score combine precision and recall into a single metric, provides a better understanding of model performance.

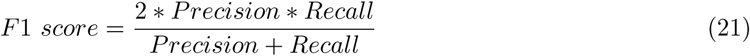

## 4 Results

### 4.1 Adjusting model parameters

#### 4.1.1 Comparison of censored normal and logistic dropout

We applied different missingness mechanisms depending on the missing proportion of each protein. When the proportion of missing values was below a threshold *λ*, we assumed that missingness arose from logistic dropout. Conversely, when the missing proportion exceeded *λ*, we modeled missingness as arising from protein-specific cutoffs. By varying *λ*, we compared the performance of the truncated normal and logistic models in terms of recall and precision. Setting *λ* = 0 implies that all missingness is attributed to cutoffs, while *λ* = 1 assumes that all missingness arises by logistic dropout. We evaluated both comparisons in the CQE dataset with known ground truth, restricting the analysis to proteins with missing values. Precision and recall were used as evaluation metrics. Overall, we observed that varying *λ* did not substantially alter model performance (Supplementary Figure S1), likely because most proteins clearly fall into one mechanism or the other, making the hybrid boundary less influential.This suggests that both missingness mechanisms capture the dominant data patterns similarly, but the hybrid framework remains useful for accommodating datasets with mixed missingness behavior.

#### 4.1.2 MCMC setting

In rjags, several parameters control the behavior of the MCMC sampler. Increasing the number of chains can improve the sensitivity of convergence diagnostics. The adaptation phase specifies the number of adaptive iterations at the beginning of a simulation, allowing the sampler to adjust proposal distributions [38]. Burn-in refers to the initial samples that are discarded because the sampler requires time to reach the stationary region of the posterior distribution. Iterations denotes the total number of steps taken by each chain (excluding burn-in). We also evaluated two strategies for initializing chains. Some recommend starting from highly dispersed initial values, under the rationale that convergence suggests full exploration of parameter space [29]. Different MCMC settings refer to Table 2. To assess the impact of these settings, we compared run time, effective sample size (ESS), Monte Carlo standard error (MCSE), accuracy, and precision under different configurations. Convergence was monitored using *R̂*, with the maximum across all parameters reported. To further assess convergence, we visually inspected trace, autocorrelation function(ACF), and posterior density plots for a representative subset of parameters, including group-level mean (*µ_jp_*) and precision 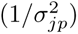. The diagnostic plots confirmed adequate mixing and approximate stationarity across chains, example figures see from Supplementary Figure S6. Each configuration was run three times, and the results were averaged. The test dataset was CQE-A, restricted to proteins with missing values (*n* = 2393).

**Table 2:** MCMC settings evaluated in rjags.

| No. | n.adapt | burn-in | n.chains | n.iter | initial |
| --- | --- | --- | --- | --- | --- |
| 1 | 500 | 500 | 2 | 5000 | centred |
| 2 | 500 | 500 | 2 | 5000 | dispersed |
| 3 | 500 | 500 | 3 | 3333 | dispersed |
| 4 | 500 | 500 | 3 | 5000 | dispersed |
| 5 | 500 | 1000 | 3 | 5000 | dispersed |
| 6 | 500 | 500 | 2 | 10000 | dispersed |
| 7 | 1000 | 500 | 2 | 10000 | dispersed |
| 8 | 500 | 500 | 4 | 10000 | dispersed |

Table 3 summarizes the influence of MCMC settings on convergence and performance metrics. Increasing the number of iterations and chains substantially improved convergence diagnostics, reflected by higher median ESS, lower MCSE, and mean *R̂* values approaching 1. An ESS of at least 1,000 is generally considered sufficient for stable posterior estimates [39], and *R̂* values below 1.01 are typically regarded as evidence of convergence [40]. While predictive performance on the CQE-A dataset (RMSE, accuracy, and precision) remained largely unchanged, convergence stability improved consistently with more extensive sampling. Parameters that failed to meet the *R̂* ≤ 1.01 criterion were mainly parameters associated with proteins containing few observations. These parameters exhibited higher autocorrelation and lower ESS, suggesting slower mixing. Applying thinning or extending the sampling period could further improve convergence for these parameters.

**Table 3:**
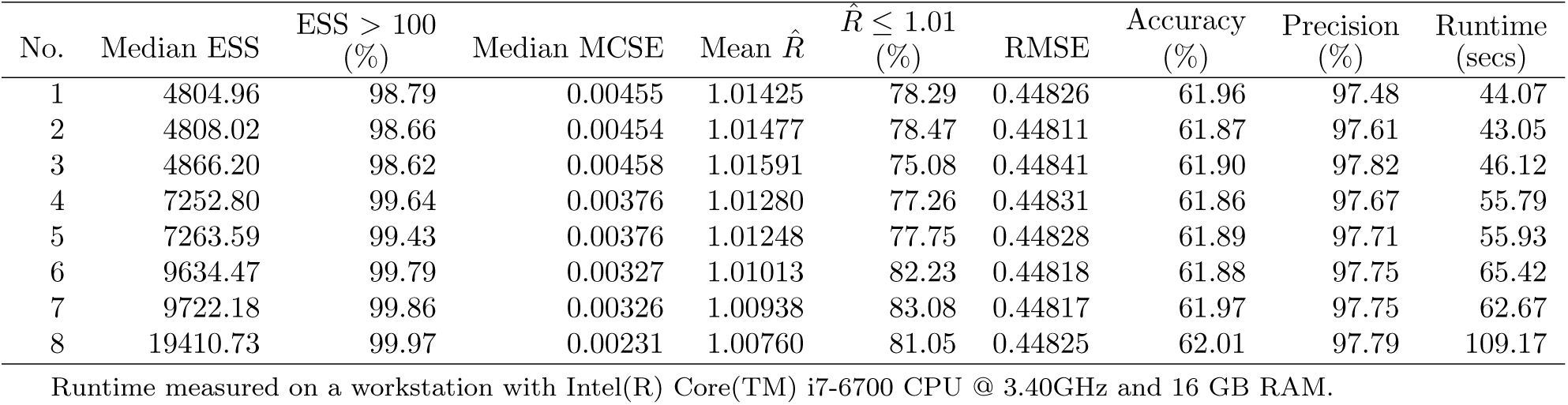
MCMC diagnostics across different settings.

Based on these observations, we selected Setting 7 as the default configuration because it achieved better overall convergence (lower *R̂*, higher ESS) without a substantial increase in runtime. In contrast, Settings 1–3 were computationally faster but exhibited weaker convergence and insufficient mixing across chains.

### 4.2 EB vs limma on model performance

As discussed in the previous section, decision rules have a substantial impact on model performance. To enable a fair comparison with `limma`, we derived z-scores from the EB model by dividing the posterior mean logFC by its posterior standard deviation and then computed two-sided p-values under a standard normal approximation. This step was performed only for comparability, since the posterior distribution of logFC is not guaranteed to be exactly normal. In `limma`, the moderated t-statistic is obtained by shrinking each sample variance toward a pooled prior estimate across all proteins [41]. Similarly, the Bayesian framework borrows strength across proteins through a mean–variance trend, where the prior on variance is modeled conditional on mean intensity. When addressing multiple comparison problems, many approaches are based on controlling false discovery rate (FDR) [32]. For `limma`, this is typically achieved using the Benjamini–Hochberg (BH) procedure [42]. For comparability, we evaluated two variants of the EB results:

1. EB-z, using p-values derived from the approximate z-scores and
2. EB-z+BH, where BH correction was applied to the p-values.

Since `limma` require group mean for each group, proteins observed in only one condition cannot be analyzed. These proteins were therefore excluded from analysis. Table 4 shows that the EB model achieves comparable performance to `limma` on CQE-A, with a slightly higher F1-score, and clearly outperforms `limma` on CQE-B, where it attains both higher precision and recall. These results suggest that the proposed EB model is robust across different data scenarios.

**Table 4:** Comparison of EB and limma on the CQE dataset.

| Dataset | Method | Precision (%) | Recall (%) | F1-Score (%) |
| --- | --- | --- | --- | --- |
| CQE-A | limma | 96.63 | 89.08 | 92.70 |
| CQE-A | EB-z+BH | 97.93 | 89.30 | 93.42 |
| CQE-A | EB-z | 95.45 | 91.63 | 93.50 |
| CQE-B | limma | 83.88 | 87.50 | 85.65 |
| CQE-B | EB-z+BH | 94.57 | 89.10 | 91.75 |
| CQE-B | EB-z | 91.32 | 92.06 | 91.69 |

We further compared the two methods in settings with no missing values, applying the full EB model. As shown in Table 5, the EB approach yields consistently higher precision and lower RMSE, particularly in the CQE-B dataset. This improvement is mainly from the limitations of the moderated *t*-test, which assumes a null hypothesis of zero effect and does not explicitly account for noise. As a result, proteins that should be negatives often appear with small but statistically significant logFC values (adj. *p <* 0.05), leading to false positives [43].We continue to observe this trend in subsequent analyses. As Andrew Gelman has argued, in practice it is rare for a true effect to be exactly zero; therefore, declaring significance solely on the basis of small *p*-values can be misleading. Our Bayesian approach mitigates this limitation by directly modeling uncertainty and noise.

**Table 5:** Comparison of EB and limma when no missing values are present.

| Dataset | Method | Precision (%) | Recall (%) | F1-Score (%) | RMSE |
| --- | --- | --- | --- | --- | --- |
| CQE-A | limma | 97.26 | 98.30 | 97.77 | 0.18667 |
| CQE-A | EB | 99.48 | 97.01 | 98.23 | 0.17623 |
| CQE-B | limma | 85.86 | 93.57 | 89.55 | 0.19125 |
| CQE-B | EB | 98.91 | 89.78 | 94.12 | 0.17353 |

### 4.3 Three species CQE dataset

To evaluate accuracy, we first applied the methods to the CQE dataset, where ground-truth fold changes are known. We defined true positives (TP) as yeast or *E. coli* proteins with the correct direction of change, and true negatives (TN) as human proteins correctly identified as not changing. We assessed model performance using Precision–Recall (PR) curves and root mean square error (RMSE). For comparison, the empirical Bayes (EB) model was evaluated alongside existing imputation methods, using default parameters. For `msImpute`, the `v2-mnar` option was selected. Differential expression (DE) analysis following imputation was performed using `limma`; “limma” alone refers to analysis without imputation.

Table 6 summarizes RMSE results and indicates whether each method supports unique proteins (i.e., proteins entirely missing in one group). The EB model achieved the lowest average RMSE across both CQE-A and CQE-B comparisons while supporting unique proteins. SVD performed second best, followed by `msImpute` and MLE, although the latter cannot process unique proteins. RMSE measures the accuracy of predicted logFC values, whereas PR curves assess a method’s ability to balance recall (recovering true positives) against precision (avoiding false positives). As shown in Figure 7, the EB model achieved the highest AUC in both comparisons, with a particularly advantage in CQE-B. This indicates improved precision–recall performance in differential expression detection, SVD remained the second strongest performer, outperforming no-imputation analysis in both settings, while `msImpute` showed robust and consistent performance. Other methods exhibited variable or unstable results.

**Figure 7:**
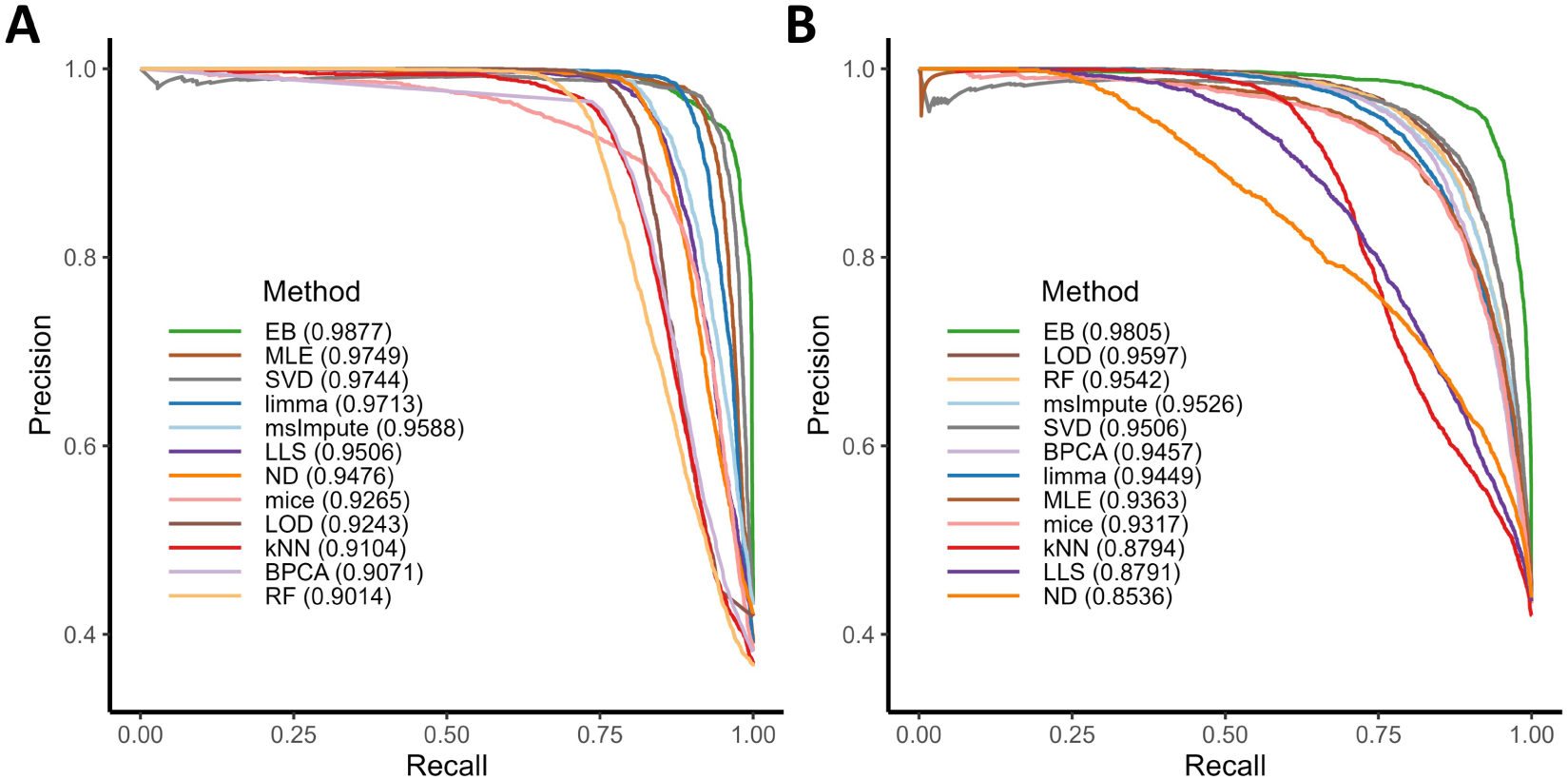
Precision–Recall curves comparing EB and existing methods under two CQE comparisons. Methods applied to (A) CQE-A and (B) CQE-B, respectively. The area under the precision–recall curve (AUPRC) is reported in the legend. *E. coli* and yeast proteins with the correct direction of change are defined as true positives and human proteins are defined as false positives.

To further visualize the predicted fold changes, we examined the distributions of differentially expressed proteins (DEPs) fold changes (yeast and *E. coli*) across methods (Figure 8). Red lines indicate the ground-truth logFC values. Ideally, a good method should yield predicted logFC values tightly centred around the ground truth. The EB model displayed the most compact distributions with minimal outliers. Notably, the EB model produced no proteins with incorrect fold-change direction in any comparison, whereas other methods occasionally produced misclassified directions.

**Figure 8:**
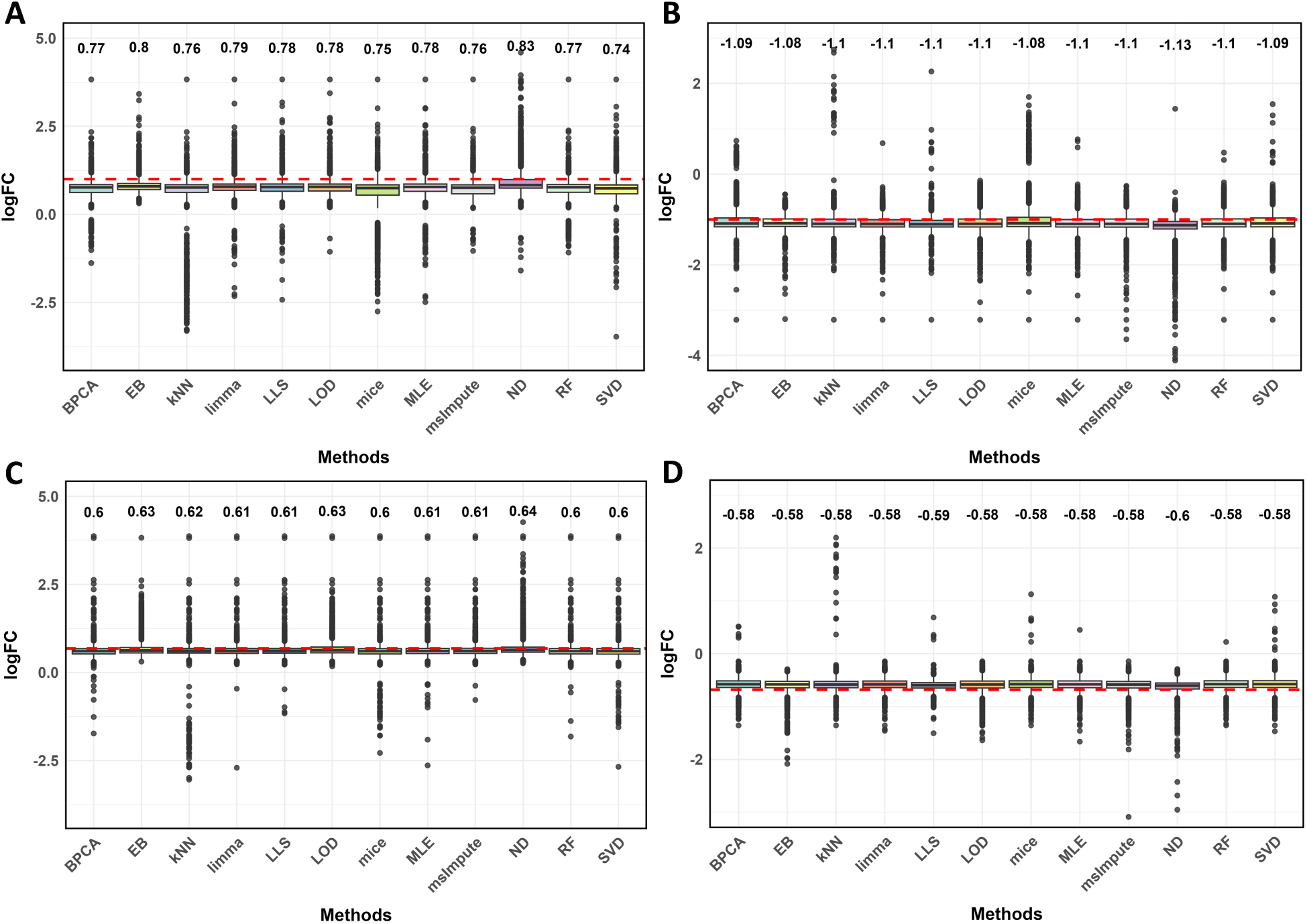
Predicted log-fold changes (logFC) for differentially expressed proteins in the CQE dataset. (A) CQE-A, yeast proteins (true logFC = 1). (B) CQE-A, *E. coli* proteins (true logFC = −1). (C) CQE-B, yeast proteins (true logFC = 0.68). (D) CQE-B, *E. coli* proteins (true logFC = −0.68). Box plots show the distribution of logFC estimates produced by each method for the corresponding protein group. Red horizontal lines indicate the true logFC values. Median logFC for each method is displayed above the respective box.

**Table 6:** Root mean square error (RMSE) comparison of tools applied to CQE dataset. “Unique proteins” indicates whether the method supports proteins with one group is entirely missing.

| Method | RMSE (A) | RMSE (B) | Unique proteins |
| --- | --- | --- | --- |
| EB | 0.2584 | 0.2413 | Y |
| SVD | 0.3026 | 0.2295 | Y |
| limma | 0.2889 | 0.2494 | N |
| msImpute | 0.3241 | 0.2151 | Y |
| MLE | 0.2963 | 0.2572 | N |
| LOD | 0.3416 | 0.2393 | Y |
| BPCA | 0.4000 | 0.2245 | Y |
| RF | 0.4175 | 0.2173 | Y |
| ND | 0.3863 | 0.3279 | Y |
| LLS | 0.3905 | 0.3410 | Y |
| mice | 0.4758 | 0.2694 | Y |
| kNN | 0.5941 | 0.3640 | Y |

We next focused on unique proteins, which are often biologically meaningful but typically excluded from conventional workflows that lack imputation. Here, we compared the EB model and imputation methods capable of handling such cases (excluding `limma` and MLE). For these proteins, the challenge lies not only in detecting differential expression but also in correctly inferring the direction of change. We therefore computed the *false sign rate*, defined as the proportion of reported positives with an incorrect direction (Type S error) [44]. As shown in Figure 9, in CQE-A the EB model achieved the lowest RMSE, 0% false sign rate, and high precision, at the expense of slightly lower recall. SVD also performed well, achieving high precision and recall but with a small (1.89%) false sign rate and slightly higher RMSE. `msImpute` exhibited higher RMSE but maintained a low false sign rate (0.35%). Other methods either had large RMSEs or high false sign rates. In CQE-B, the EB model remained competitive, with a 0% false sign rate and balanced precision–recall performance. `msImpute` achieved the best overall trade-off, with the lowest RMSE and strong recall, while SVD showed unstable behavior with a marked increase in false sign rate. RF and ND were overly conservative, missing many true signals.

**Figure 9:**
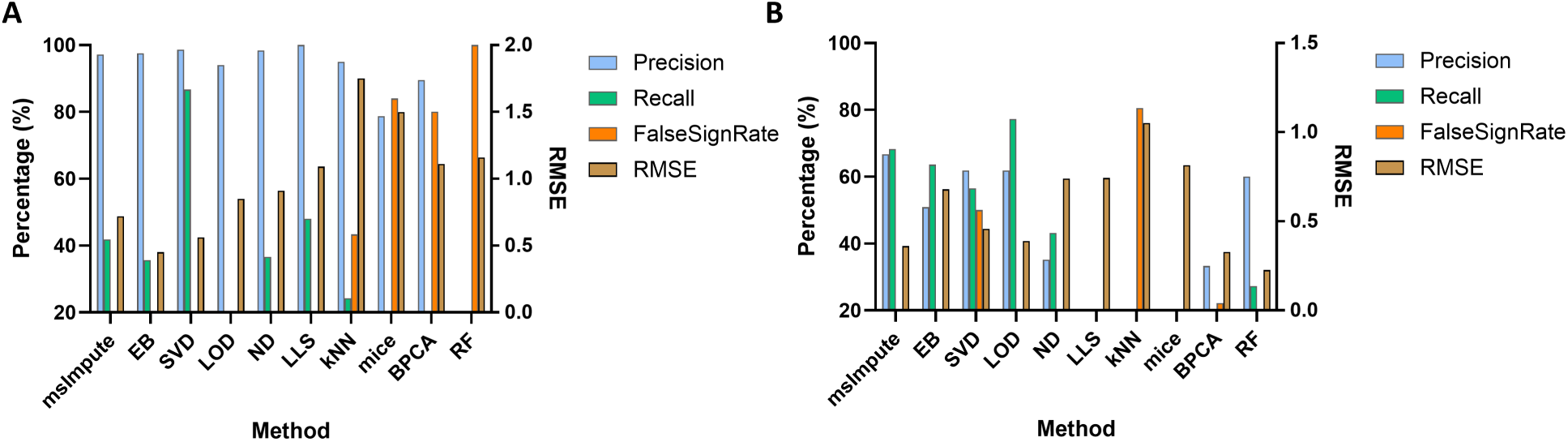
Performance comparison on unique proteins for diffent methods applied to the CQE dataset. (A) Precision, recall, False sign rate and RMSE for CQE-A. (B) Precision, recall, False sign rate and RMSE for CQE-B.

Overall, we conclude that for unique proteins, the EB model provides the most reliable and stable performance, consistently achieving a 0% false sign rate across datasets. It is conservative yet robust. `msImpute` serves as a strong alternative, offering a more balanced trade-off between precision and recall, whereas other methods tend to be less stable or more error-prone.

### 4.4 phagoFACS dataset

While the CQE dataset serves as a useful benchmark for method evaluation, it does not fully capture the biological variability observed in real experiments. To assess our EB model under realistic conditions, we benchmarked it using the phagoFACS dataset derived from macrophages infected with STM. To construct a ground-truth reference, we first removed all proteins with missing values, resulting in a complete dataset where all intensities were known. We then introduced missingness artificially under two mechanisms and at varying proportions (5%, 10%, 20%, and 30%). In the first mechanism, missingness followed a MNAR pattern dependent on intensity, generated via logistic functions. The EB model explicitly accounts for MNAR missingness through its logistic-missingness framework, with the threshold parameter set to 1. The complete dataset was analyzed using a standard *t*-test to obtain “true” logFC and *p*-values, with proteins having *p <* 0.05 labeled as true positives. To test robustness beyond the logistic MNAR assumption, we also introduced an alternative missingness scenario using protein-specific cutoffs sampled from normal distributions, setting values below each cutoff to missing (NA) and adding an additional 1% MCAR noise. The distributions before filtering, before introducing missing values, and after introducing missing values (supplementary Figure S2 and S3) confirmed that the dataset with certain values set to missing retained similar distributional shapes to the original, ensuring that the benchmarks remain representative. We evaluated all methods using P-R curves, RMSE, and predictive fit.

#### Logistic MNAR missingness

Under the logistic MNAR setting (Figure 10), the EB model consistently achieved high AUC across all missingness levels, followed by no-imputation and MLE. `msImpute`, LOD, and ND also performed well, showing stable results just behind the top performers. In contrast, SVD, which was strong on the CQE dataset, performed poorly in this setting. In terms of RMSE (Table 7), the EB model ranked third-lowest, only behind `limma` and MLE. However, it should be noted that these two methods exclude unique proteins, which artificially lowers their RMSE values.

**Figure 10:**
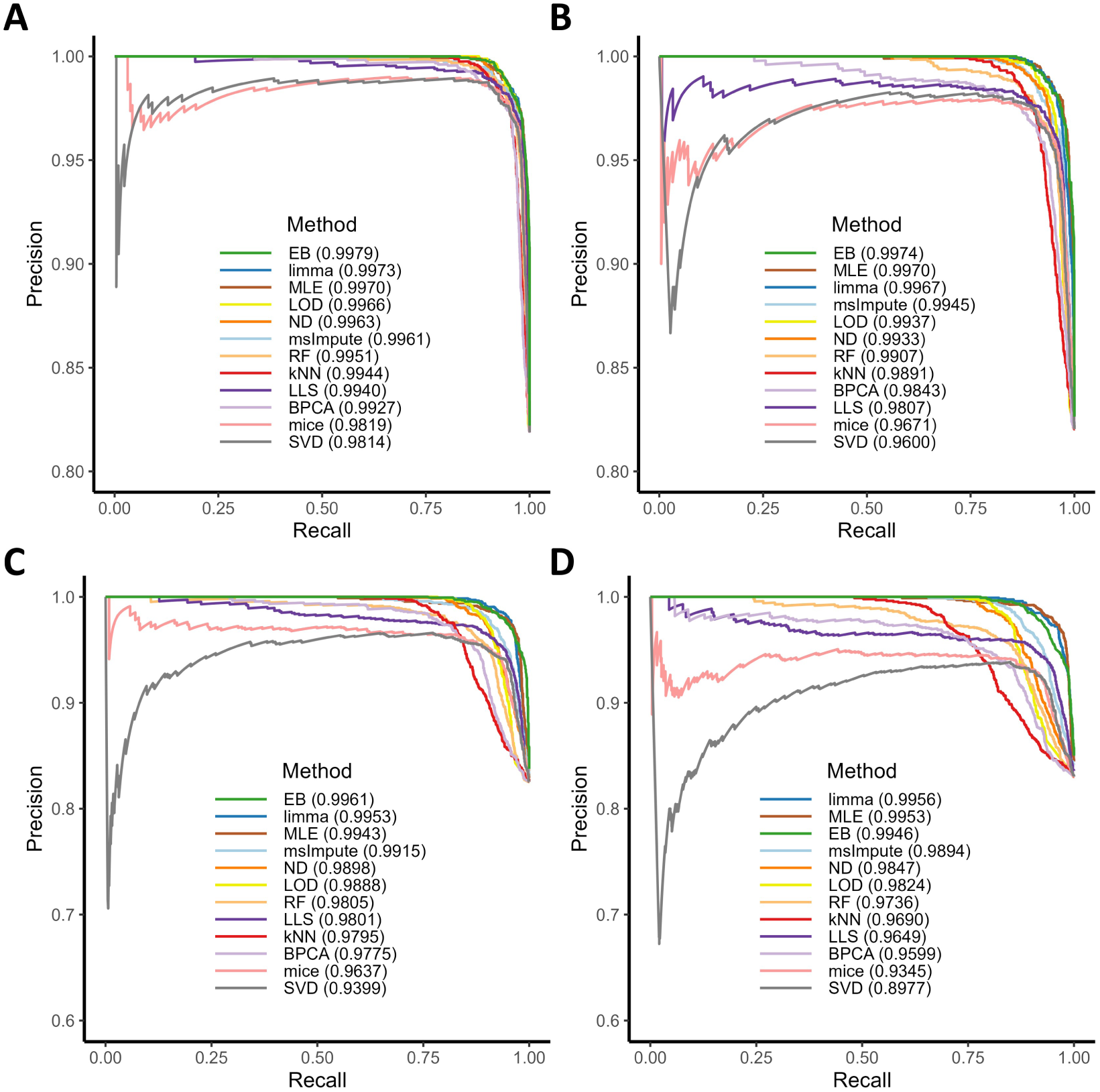
Precision–Recall curves under logistic MNAR missingness at varying missingness levels. (A) 5% missingness. (B) 10% missingness. (C) 20% missingness. (D) 30% missingness. The area under the precision–recall curve (AUPRC) is reported in the legend. True positives are defined as proteins with *p <* 0.05 under the standard *t*-test, and false positives as proteins with *p* ≥ 0.05.

**Table 7:** RMSE of methods under logistic MNAR missingness at varying proportions. *Methods that exclude unique proteins.

| Method | 5% | 10% | 20% | 30% |
| --- | --- | --- | --- | --- |
| limma* | 0.1169 | 0.1327 | 0.1537 | 0.1701 |
| MLE* | 0.1228 | 0.1477 | 0.1675 | 0.1853 |
| EB | 0.1235 | 0.1533 | 0.2111 | 0.2784 |
| ND | 0.1396 | 0.2053 | 0.2920 | 0.3972 |
| msImpute | 0.1703 | 0.2450 | 0.3354 | 0.3868 |
| LOD | 0.1351 | 0.2163 | 0.3572 | 0.4789 |
| RF | 0.2165 | 0.3327 | 0.4706 | 0.6008 |
| BPCA | 0.2667 | 0.3915 | 0.5471 | 0.6542 |
| mice | 0.3586 | 0.4810 | 0.6576 | 0.7560 |
| SVD | 0.2695 | 0.4595 | 0.7306 | 0.9075 |
| kNN | 0.3192 | 0.4982 | 0.7391 | 0.8584 |
| LLS | 0.2187 | 1.0893 | 1.0203 | 1.1289 |

To further assess imputation quality, we compared imputed distributions against the original data (Figure 11). Since the EB model does not generate explicit imputed values, we instead sampled draws from the posterior predictive distribution to construct the predictive fit. The EB model and ND most closely reproduced the original distribution, while `msImpute` also performed well. Other methods distorted the shape of the distribution, often overestimating mid-range values and underestimating low-intensity ones.

**Figure 11:**
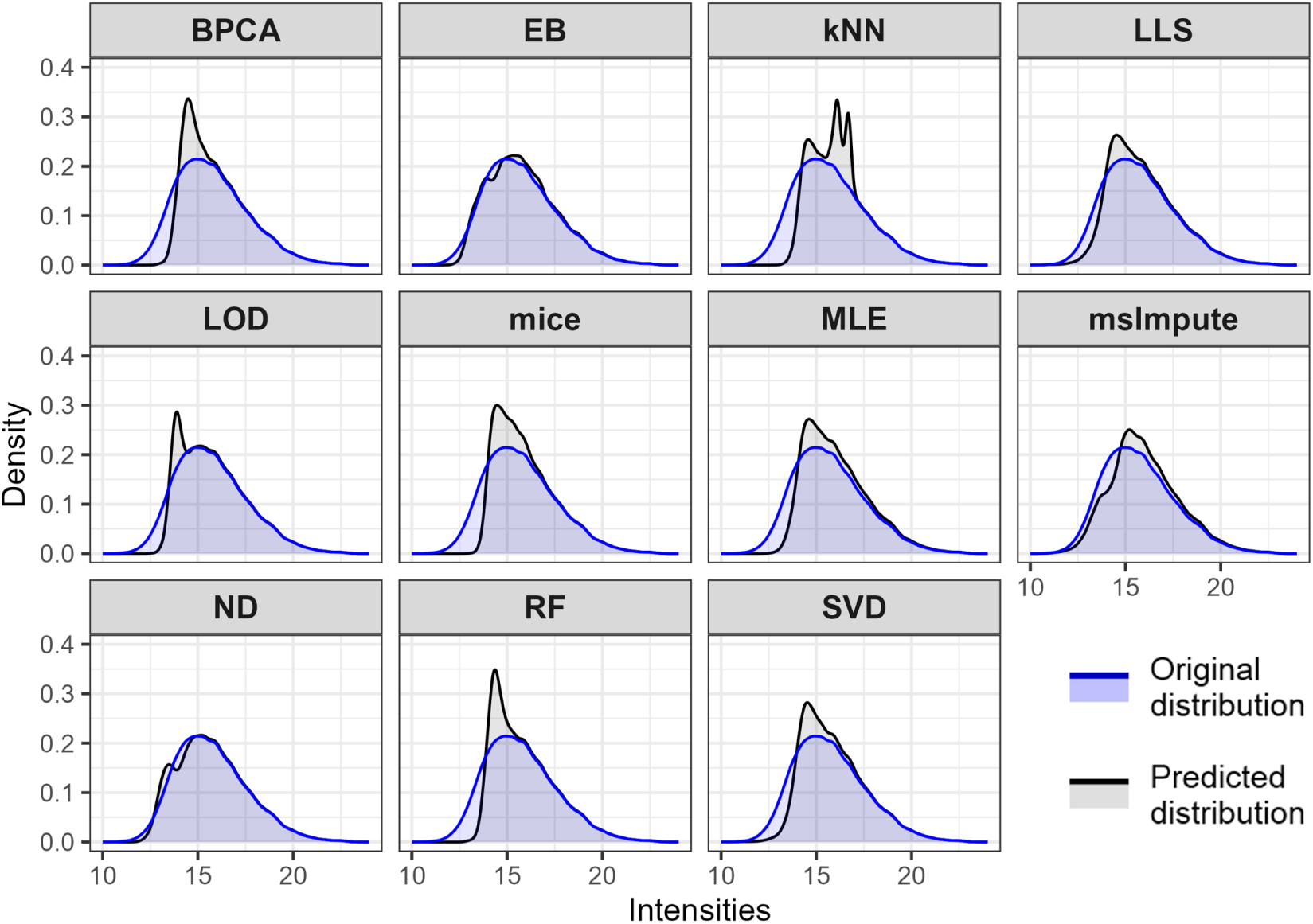
Predictive fit for a dataset with 20% values artificially set to missing under a logistic MNAR mechanism. The blue curve shows the distribution of the original (complete) data, and the black curves show the distributions predicted by each method. For EB, predicted values were generated from the posterior predictive distribution.

#### Protein-specific cutoff + MCAR missingness

We next evaluated performance under missingness generated from protein-specific cutoffs (sampled from a normal distribution) with an added 1% MCAR noise. As before, four missingness levels were tested. Across all levels, the EB model again showed strong and stable performance, ranking third in RMSE after `limma` and MLE (Supplementary Table S2). In PR analysis (Supplementary Figure S4), the EB model achieved similar AUC with no-imputation and MLE. LOD, and `msImpute` also performed well, maintaining low RMSE and stable behavior across missingness levels. In predictive-fit evaluation (Supplementary Figure S5), EB, `msImpute`, and ND best preserved the original data distribution, whereas other methods exhibited clear deviations or bias in specific intensity ranges.

Across both missingness mechanisms, the EB model consistently demonstrated robust and reliable performance. It achieved the best overall PR AUCs and competitive RMSE values while preserving realistic data distributions. Notably, `limma` also performed strongly, suggesting that imputation can sometimes distort biological data, as seen in predictive-fit comparisons. When imputation is necessary, MLE, `msImpute` and ND provided the most accurate and stable alternatives.

### 4.5 UPS spike-in dataset

Next, we evaluated the false discovery rate (FDR) control of different methods using the UPS spike-in dataset. This dataset is particularly useful because the vast majority of proteins remain unchanged, while only 48 spiked-in proteins differ across groups. Thus, it provides a clean benchmark for assessing Type I error control. We compared nine spike-in concentrations against the 500 amol baseline (Table 8 and Figure 12). Performance was evaluated in terms of mean true positive rate (TPR), mean precision (1–FDR), and the range of observed precision values across comparisons. Across all nine spike-in comparisons, the EB model maintained a relatively stable precision (approximately 40–80%), while other methods exhibited wide fluctuations (e.g., `limma`: 5–100%). Importantly, this stability was preserved even at higher fold-changes (10–50 fmol), where competing methods tended to substantially inflate false discoveries. Although the reported precision appeared similar across methods, the absolute number of false positives varied widely. For example, in the 50 fmol vs 500 fmol comparison, the EB model reported only 73 false positives, whereas most other methods reported close to 1000 — inflating the discovery set dramatically. These results demonstrate that the Bayesian framework provides more reliable control of false discoveries, maintaining balanced sensitivity and precision across a wide range of spike-in concentrations.

**Figure 12:**
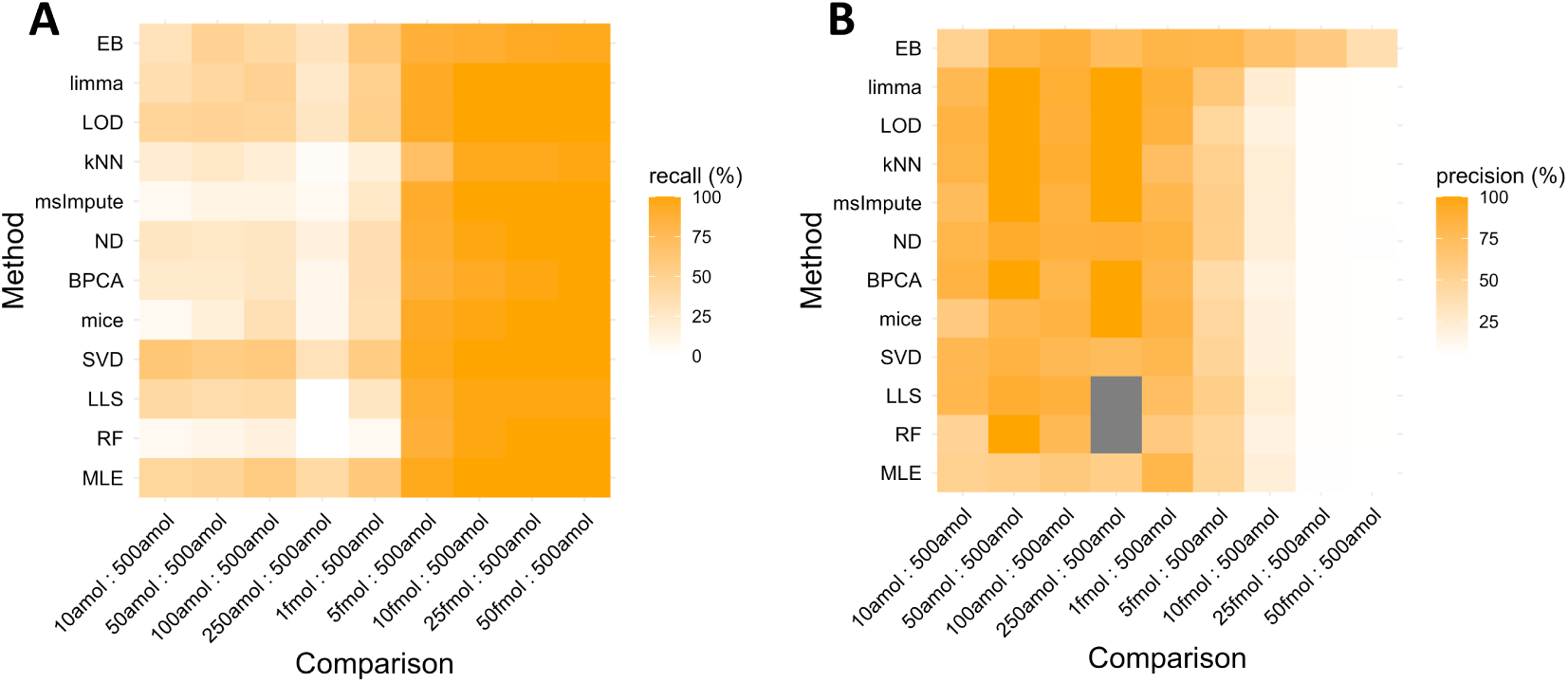
Performance of different methods across nine comparisons in the UPS spike-in dataset. Heatmaps show (A) recall and (B) precision, with color intensity indicating performance (higher values correspond to better performance). True positives are defined as 48 spiked proteins. Grey squares indicate cases in which a method detected no positive proteins.

**Table 8:** Summary performance of different methods across nine UPS spike-in comparisons.

| Method | Mean Recall (%) | Mean Precision (%) | Min Precision (%) | Max Precision (%) |
| --- | --- | --- | --- | --- |
| EB | 64.56 | 69.45 | 38.66 | 86.36 |
| limma | 66.18 | 61.44 | 4.49 | 100 |
| LOD | 68.06 | 59.35 | 4.68 | 100 |
| kNN | 49.54 | 58.97 | 4.69 | 100 |
| msImpute | 50.46 | 58.58 | 4.63 | 100 |
| ND | 57.41 | 57.58 | 4.77 | 92.31 |
| BPCA | 55.09 | 57.14 | 4.47 | 100 |
| mice | 54.40 | 53.74 | 4.49 | 100 |
| SVD | 73.38 | 52.57 | 4.47 | 84.38 |
| LLS | 58.80 | 52.12 | 4.56 | 90.00 |
| RF | 46.99 | 45.39 | 4.58 | 100 |
| MLE | 71.67 | 42.67 | 4.25 | 81.82 |

### 4.6 Enrichment analysis

Following DE analysis, enrichment analysis is often performed to identify the biological processes associated with the observed changes in protein abundance. We evaluated the performance of different methods in pathway analysis using an immune cell dataset containing 10,465 proteins across 175 samples, following the design of Jin et al. [7].

For `msImpute`, RF, PCA methods, only two groups under comparison were retained for imputation, as these methods did not converge within a reasonable time. For each immune cell type, DEPs between the activated and steady-state conditions were identified using BH-adjusted *p*-values *<* 0.05, or, for the EB model, using posterior probabilities *p_>ROP E_* or *p_<ROP E_ >* 0.95. Enriched Gene Ontology (GO) biological processes were then determined from the DEPs.

We focused on pathways biologically relevant to cell activation. Specifically, for B cells, we searched for terms containing “B cell” or “lymphocyte activation”; for T4 cells, we used keywords such as “T cell activation,” “T lymphocyte activation,” “CD4,” “helper T,” “T cell receptor,” “TCR signaling,” and “T cell–mediated immunity.” We summarized the number of enriched pathways, total genes, and pathways with adjusted *p*-values *<* 0.05 for each method in Supplementary Table S3 and S4.

Overall, the EB, LLS, and mice methods identified the largest number of significant pathways related to B cell activation, while SVD, mice, and LLS performed best in T4 cell activation. In B cell activation–related pathways, the EB model, LLS, and SVD showed stronger enrichment signals. In contrast, for T4 cell activation, SVD, LLS, and mice achieved the best performance, followed by EB and ND. In both cell types, omitting imputation markedly reduced the number of significant pathways detected, highlighting the importance of imputation for meaningful downstream biological interpretation. These results suggested that the EB model provides robust performance for enrichment analysis.

**Figure 13:**
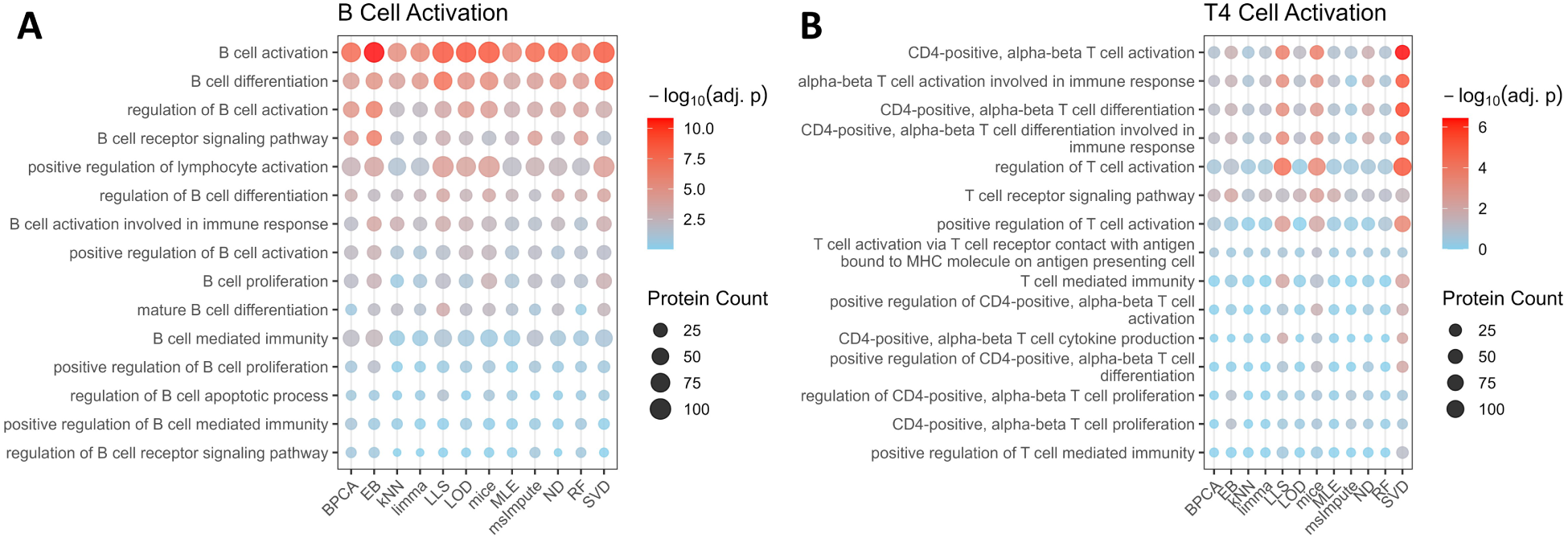
Pathway enrichment analysis in the immune dataset. Panels show selected pathways enriched among differentially expressed proteins associated with (A) B cell activation and (B) T4 cell activation. Pathways were chosen based on their relevance to the corresponding biological process.

## 5 Discussion

The presence of missing values is a key challenge in label-free proteomics studies. Differential expression analysis methods that explicitly account for missingness can improve proteome coverage and increase statistical power. In this study, we demonstrated that the EB model outperforms existing imputation-based methods as well as the benchmark method `limma`, providing a more comprehensive view of protein-level changes between conditions.

We investigated model performance under different hypotheses of missingness. Whether missing values arose from protein-specific cutoffs or from a logistic probability related to latent intensity, the results were broadly similar in terms of recall and precision, suggesting that the underlying missingness mechanism is complex. We also assessed the influence of MCMC settings and found that the dispersion of initial values, the number of adaptation steps, and the length of burn-in had little impact on model convergence, diagnostics, or performance. Increasing the total number of sampling steps improved effective sample size, MCSE, and *R̂*, but at the cost of longer run time, without a clear gain in performance. Based on these findings, we recommend starting with fewer or shorter chains, evaluating convergence with standard diagnostics, and then adjusting settings only if necessary.

We compared the EB model with `limma` using the same decision rule. Across two comparisons, the EB model consistently achieved a higher F1-score, regardless of whether FDR correction was applied. While the Bayesian and frequentist frameworks are not directly comparable, these results provide a useful perspective for disentangling the contribution of modelling (how fold changes and uncertainties are estimated) from significance testing (how differential abundance is declared). Importantly, they demonstrate that the EB model remains robust even when evaluated under a frequentist decision rule. For proteins without missing values, the EB model achieved higher precision at the expense of recall. Nevertheless, the overall F1-score was higher and the RMSE was lower, indicating improved accuracy even in the absence of missingness.

From the CQE dataset, we found that the EB model achieved the highest AUC in both precision–recall curves, while also yielding the smallest average RMSE among all methods. We also showed that not all imputation strategies evaluated here are capable of handling unique proteins, highlighting the need for caution when selecting appropriate imputation methods. Together, these results indicate that the EB model offers both strong precision and accurate logFC estimation. Using a real proteomics dataset with artificially introduced missing values, we further demonstrated that the EB model consistently outperformed other methods across four levels of missingness, in terms of both higher AUC and lower RMSE. The exceptions were `limma` and MLE, which sometimes achieved lower RMSE, likely because they do not account for unique proteins. We also compared predictive fits against the complete dataset and found that most imputation methods failed to recover the original data distribution. By contrast, the EB model, `msImpute` and ND provided the best fits. Because lower FDR (i.e. higher precision) is critical in proteomics data analysis, we also evaluated performance using the UPS spike-in dataset, where only 48 proteins were differentially expressed. Here, the EB model consistently achieved moderate recall across all comparisons. In the nine comparisons, it maintained moderate precision, whereas most other methods exhibited extremely low precision in at least some comparisons. This suggests that the EB model offers more stable performance across diverse scenarios.

Lastly, we evaluated the performance of the EB model in pathway enrichment analysis using an immune dataset. The results showed that the EB model performed particularly well for B cell activation, ranking closely behind SVD, LLS, and mice.

In summary, the EB model provides a powerful framework for differential expression analysis in proteomics. It offers robust performance, high accuracy, and explicit quantification of uncertainty, while eliminating the need for imputation and making effective use of information contained in missingness patterns. By presenting a complete Bayesian workflow—from decision making to result interpretation—we aim to illustrate how Bayesian methods can be effectively applied in proteomics, a field still largely dominated by frequentist approaches, and to encourage their broader adoption. Looking forward, further extensions could involve implementing the model for single-cell proteomics, where missing values are abundant, and systematically evaluating the power of Bayesian analysis across varying levels of missingness/sample sizes. Moreover, the lack of consensus on the definition and choice of the ROPE warrants further discussion within the community to enhance the interpretability and comparability of Bayesian results in proteomics research.

## 6 Data and Code availability

The proteomics datasets analyzed during this study are publicly available in the PRIDE repository under identifiers PXD062621, PXD068192, PXD072249, PXD009815 and PXD004352.

Our implementation is provided as the R package `MissBayes`, which is currently available on GitHub at: https://github.com/lmcdbd/MissBayes. The package supports multiple input formats including log-intensity matrices, data frames, and `QFeatures` objects. Source code and all analysis scripts used in this study are available at: https://github.com/lmcdbd/MissBayes-DataAnalysis.

## 7 Conflict of interest

The authors declare that they have no known competing financial interests or personal relationships that could have appeared to influence the work reported in this paper.

## Acknowledgments

ML is being funded by a EPSRC studentship co-funded by GSK within the MosMed Centre for Doctoral Training. The work was partly funded by a Wellcome Investigator Award (215542/Z/19/Z) to MT and a Wellcome Trust multi-user equipment grant (212947/Z/18/Z).

## Notes

### Competing Interest Statement

The authors have declared no competing interest.

