## Supplementary material for "Empirical-Bayes and Bayesian Hierarchical Modelling for Missingness and Differential Expression in Proteomics": supplementary (5).pdf

**Contents**

- Section S1: Proteomics data aqcuisition and processing
- Section S2: Calculation of the parameters of the Inverse-Gamma prior
- Table S1: Summary of datasets
- Table S2: RMSE of methods under cutoff and MCAR missingness at varying proportions
- Table S3: Summary of enrichment analysis (B cells)
- Table S4: Summary of enrichment analysis (T4 cells)
- Figure S1: Comparison of censored normal and logistic dropout
- Figure S2: Data distribution under logistic MNAR
- Figure S3: Data distribution under cutoffs and MCAR
- Figure S4: Precision–Recall curves under cutoff and MCAR missingness at varying proportions
- Figure S5: Predictive fit of dataset with 20% values set to missing under cutoff and MCAR missingness
- Figure S6: Example of MCMC diagnostic plots

#### S1. Proteomics data acquisition and processing

All proteomics data acquisition was performed as part of previous work with the methods and mass spectrometry datasets publicly available, but the pooled liver dataset was not reported in its respective study [18]. Thus, here we describe its acquisition.

Samples were derived from pooled liver biopsies and underwent processing to tryptic peptides as described previously [18]. Immediately before acquisition, samples were resuspended in 0.1 % formic acid and a volume equivalent to 500 ng of starting material was subjected to TIMS-LC-MS using a NanoElute II LC in line with a timsToF-HT (Bruker). For LC, two-column separation was used, with a ThermoFisher 5mm Trap Cartridge and a Bruker FIFTEEN (15cm) separation column at 50°C, and a 20 µm captivespray emitter. Buffer A was 0.1 % formic acid in water, buffer B was 0.1 % formic acid in acetonitrile. Equilibration was performed with 10 x column volumes for trap and 4 column washes for the elution columns. Three different gradient lengths were used: Buffer B was increased from 5 % to 35 % for 60, 120, or 180 minutes, with flow rate maintained at 300 nl/min. Then all three gradients were proceeded by increasing buffer B to 95 % and flow rate to 400 nl/min in the next 30 seconds and holding for 3 minutes, then increasing the flow rate to 600 nl/min over the next 2 minutes. The mass spectrometer was operated in ddaPASEF mode. Mass and IM ranges were 100-1700 m/z and 0.6-1.45 1/K0. TIMS ramp and accumulation times were 100 ms, with a total of 10 PASEF ramps, with a target intensity of 20000 and threshold of 2500. Total cycle time was 1.17 seconds . Collision energy was applied in a linear fashion based on ion mobility, where ion mobility = 0.6-1.6 1/K0, and collision energy = 20-59 eV. ddaPASEF data files were searched using FragPipe v20.0 [16], with the built in LFQ-default pipeline. The proteome FASTA file was the reference proteome for Homo sapiens (UP000001414 SwissProt, with isoforms, downloaded from UniProt on the 10th of March 2023) combined with a common contaminants list [45]. MSFragger settings included precursor and fragment mass tolerance of ±20, cysteine carbamidomethylation set as fixed, oxidation of methionine, protein N-terminal acetylation as variable modifications, trypsin specificity with 2 missed cleavages were permitted, peptide length 7-50, mass 500-5000 Da. Percolator was used for PSM validation, with peptide probability set to 0.5. Quantification was performed with IonQuant [46,47]. MBR was not performed, no cross run normalisation was performed at this stage.

For the pooled QC dataset, raw diaPASEF data files were searched using DIA-NN V 1.9 [17], using its in silico generated spectral library function, based on reference proteome FASTA files for H. sapiens (UP000001414, downloaded from UniProt on 11/05/2023) and a common contaminants list [45]. Trypsin specificity with a maximum of 1 missed cleavage was permitted per peptide, cysteine carbamidomethylation were set as a fixed modification. Peptide length and m/z was 7-30 and 300-1200, charge states 2-4 were included. Mass accuracy was fixed to 15 ppm for MS1 and MS2. Protein and peptide FDR were both set to 1 %. All other settings were left as defaults.

#### S2. Calculation of the Parameters of the Inverse-Gamma Prior

Assume that

$$\sigma_{jp}^2 \sim Inv - Gamma(\alpha, \beta),$$

For each group $j$, protein $p$,

$$s_{jp}^2 = \frac{1}{n-1} \sum_{1 \leq i \leq n} (y_{ijp} - \bar{y}_{jp})$$

is the sample variance, where $n$ is the number of samples per group.
Bin sample means into intervals with the width of 1, given:

$$\overline{x} = \text{sample mean of Vars}$$
$$s^2 = \text{sample variance of Vars}$$

Then

$$E[s_{jp}^2 | \sigma_{jp}^2] = \sigma_{jp}^2.$$

Therefore

$$\overline{x} = E[s_{jp}^2] = E[E[s_{jp}^2 | \sigma_{jp}^2]] = E[\sigma_{jp}^2].$$

We now calculate

$$E[s^2] = \text{Var}(s_{jp}^2) = E[s_{jp}^4] - (E[s_{jp}^2])^2 = E[E[s_{jp}^4 | \sigma_{jp}]] - (E[\sigma_{jp}^2])^2.$$

We see that

$$E[s_{jp}^4 | \sigma_{jp}] = \frac{n}{n-1}(1 - \frac{1}{n^2})\sigma_{jp}^4 = \frac{n+1}{n}\sigma_{jp}^4.$$

Therefore

$$E[s^2] = \frac{1}{n} E[\sigma_{jp}^4]$$

We therefore summarize this as

$$E[\overline{x}] = E[\sigma_{jp}^2], E[s^2] = \frac{1}{n} E[\sigma_{jp}^4].$$

We see that
$$E[\sigma_{jp}^2] = \frac{\beta}{\alpha - 1}, \quad E[\sigma_{jp}^4] = \frac{\beta^2}{(\alpha-1)^2(\alpha-2)}.$$
Therefore we should take $\alpha$ and $\beta$ satisfying
$$\bar{x} = \frac{\beta}{\alpha - 1}, \quad s^2 = \frac{\beta^2}{n(\alpha-1)^2(\alpha-2)}.$$
We now solve the algebra. We see that $\beta = (\alpha - 1)\bar{x}$ so
$$s^2 = \frac{\bar{x}^2}{n(\alpha - 2)},$$
hence
$$\alpha = \frac{\bar{x}^2}{ns^2} + 2,$$
This implies $\beta = \bar{x} \cdot (\frac{\bar{x}^2}{ns^2} + 1)$. We summarize this as
$$\beta = \frac{\bar{x}^3}{ns^2} + \bar{x}, \quad \alpha = \frac{\bar{x}^2}{ns^2} + 2.$$

#### Table S1: Summary of datasets

| **Dataset** | **Comparisons investigated** | **Number of proteins** | **Runs/Groups/ Missing proportion (%)** | **Ground truth (logFC)** |
|---|---|---|---|---|
| Pooled liver samples | NA | 3973 | 3/4/21.4 | NA |
| Pooled QC samples | NA | 7705 | 11/1/1.1 | NA |
| CQE | HYE70_10_20:HYE70_20_10 (A) | human: 6400 | 5/2/8.8 | 0 |
| | | E.coli: 2132 | | -1 |
| | | YEAST: 2832 | | 1 |
| | HYE35_20_45:HYE35_45_20 (B) | human: 6265 | 5/2/7.8 | 0 |
| | | E.coli: 2154 | | -0.68 |
| | | YEAST: 2930 | | 0.68 |
| phagoFACS | PhoP_4h: PhoP_UP | 5689 | 6/6/18.2 | NA |
| UPS spike-in | 10amol:500amol | UPS: 48; Yeast: 2729 | 4/2/6.9 | UPS: -5.6; Yeast: 0 |
| | 50amol:500amol | | 4/2/6.6 | UPS: 3.3; Yeast: 0 |
| | 100amol:500amol | | 4/2/6.7 | UPS: 2.3; Yeast: 0 |
| | 250amol:500amol | | 4/2/6.4 | UPS: 1; Yeast: 0 |
| | 1fmol:500amol | | 4/2/6.1 | UPS: -1; Yeast: 0 |
| | 5fmol:500amol | | 4/2/6.0 | UPS: -3.3; Yeast: 0 |
| | 10fmol:500amol | | 4/2/6.0 | UPS: -4.3; Yeast: 0 |
| | 25fmol:500amol | | 4/2/6.2 | UPS: -5.6; Yeast: 0 |
| | 50fmol:500amol | | 4/2/6.5 | UPS: -6.6; Yeast: 0 |
| Immune | B_naive_activated: B_steady_state | 9351 | 4/2/30.3 | NA |
| | T4_naive_activated: T4_steady_state | 9169 | 4/2/30.0 | NA |

**Table S1.** Summary of datasets used in this study.

#### Table S2: RMSE of methods under cutoff and MCAR missingness at varying proportions.

| Method | 5% | 10% | 20% | 30% |
|---|---|---|---|---|
| limma* | 0.1160 | 0.1451 | 0.1544 | 0.1747 |
| MLE* | 0.1255 | 0.1471 | 0.1587 | 0.1770 |
| EB | 0.1845 | 0.2134 | 0.2739 | 0.3720 |
| msImpute | 0.1711 | 0.2332 | 0.3287 | 0.3957 |
| ND | 0.2483 | 0.2768 | 0.3284 | 0.4079 |
| LOD | 0.1743 | 0.2721 | 0.3960 | 0.5312 |
| RF | 0.2298 | 0.3615 | 0.5088 | 0.6510 |
| bpca | 0.2572 | 0.4075 | 0.5544 | 0.6922 |
| SVD | 0.2661 | 0.4611 | 0.6537 | 0.8489 |
| knn | 0.3097 | 0.4714 | 0.6675 | 0.8346 |
| mice | 0.3436 | 0.5470 | 0.7178 | 0.8460 |
| LLS | 0.2434 | 0.4399 | 0.7342 | 1.4367 |

**Table S2.** RMSE of methods under cutoff and MCAR missingness at varying proportions. *Methods that exclude unique proteins.

#### Table S3: Summary of enrichment analysis results (B cells)

| **Method** | **Number of pathways** | **Number of genes** | **Number of pathways (adj. $p$ value $< 0.05$)** |
|---|---|---|---|
| EB | 30 | 661 | 20 |
| LLS | 29 | 789 | 20 |
| mice | 29 | 784 | 18 |
| SVD | 29 | 758 | 17 |
| bpca | 30 | 587 | 15 |
| ND | 29 | 519 | 13 |
| RF | 30 | 485 | 13 |
| MLE | 28 | 571 | 12 |
| knn | 29 | 480 | 12 |
| LOD | 29 | 612 | 12 |
| msImpute | 30 | 548 | 11 |
| limma | 29 | 497 | 10 |

**Table S3.** Summary of enrichment analysis results across different imputation or modeling methods (B cells). The last column shows the number of significantly enriched pathways with adjusted $p$ value $< 0.05$.

#### Table S4: Summary of enrichment analysis results (T4 cells)

| Method | Number of pathways | Number of genes | Number of pathways (adj. $p$ value $< 0.05$) |
|---|---|---|---|
| SVD | 42 | 947 | 23 |
| mice | 42 | 818 | 18 |
| LLS | 42 | 763 | 15 |
| ND | 42 | 475 | 11 |
| EB | 41 | 521 | 7 |
| LOD | 42 | 471 | 4 |
| bpca | 42 | 354 | 2 |
| MLE | 42 | 478 | 2 |
| limma | 42 | 415 | 2 |
| RF | 42 | 410 | 1 |
| knn | 42 | 386 | 0 |
| msImpute | 42 | 416 | 0 |

**Table S4.** Summary of enrichment analysis results across different imputation or modeling methods (T4 cells). The last column shows the number of significantly enriched pathways with adjusted $p$ value $< 0.05$.

#### Figure S1: Comparison of censored normal and logistic dropout

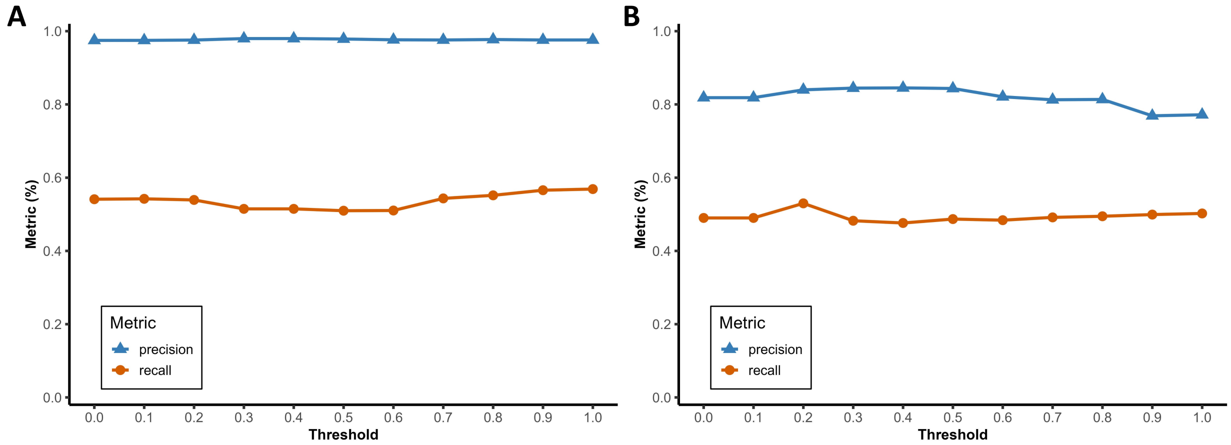


**Figure S1.** Comparison of the truncated normal and logistic dropout mechanisms across different $\lambda$ thresholds in the CQE datasets. Threshold $\lambda$ controls whether missingness is modeled as censoring or random dropout. Both mechanisms yield similar precision and recall, suggesting that the choice of $\lambda$ has limited impact on overall performance.(A) CQE-A. (B) CQE-B.

#### Figure S2: Data distribution under logistic MNAR

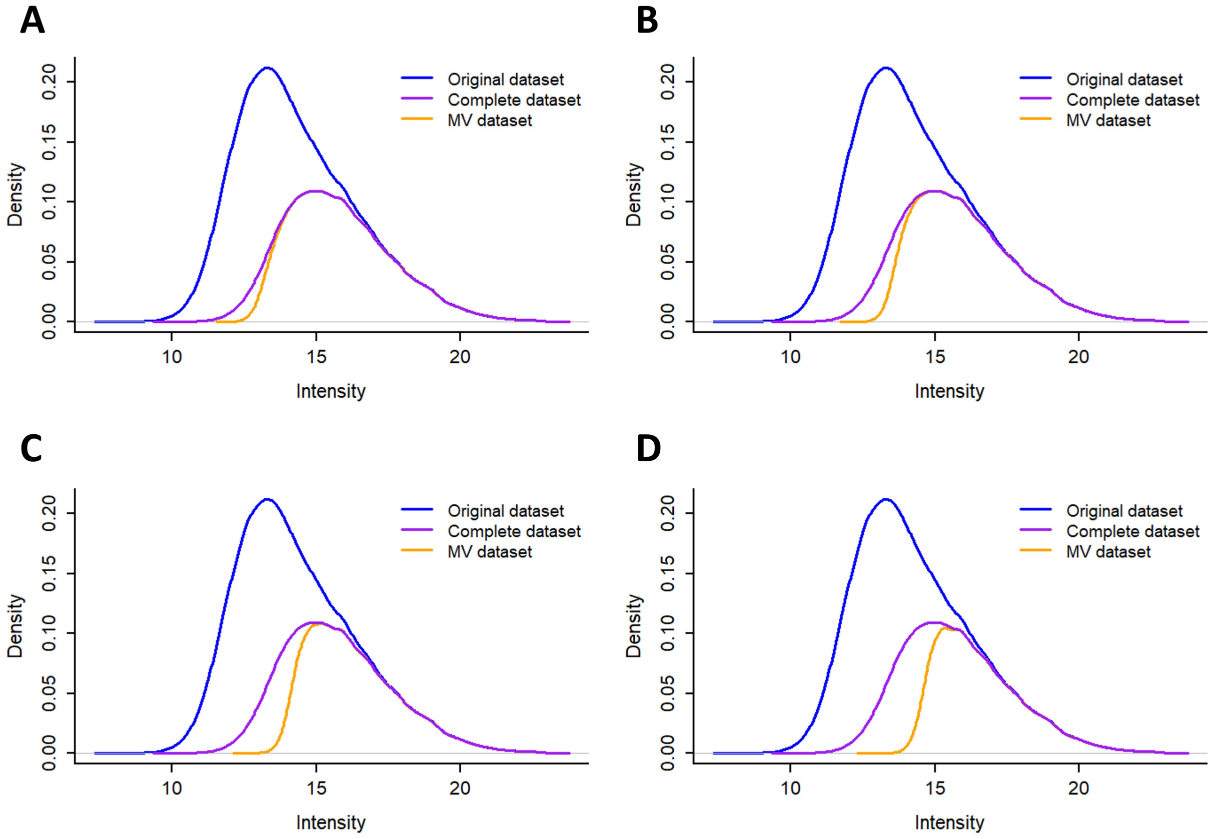


**Figure S2.** Data distribution under logistic MNAR at missingness levels of (a) 5%, (b) 10%, (c) 20%, and (d) 30%. The blue curve shows the original distribution with missing values; the purple curve shows the distribution after removing proteins with missing values; and the yellow curve shows the distribution after introducing logistic MNAR missingness into the complete dataset.

### Figure S3: Data distribution under cutoffs and MCAR

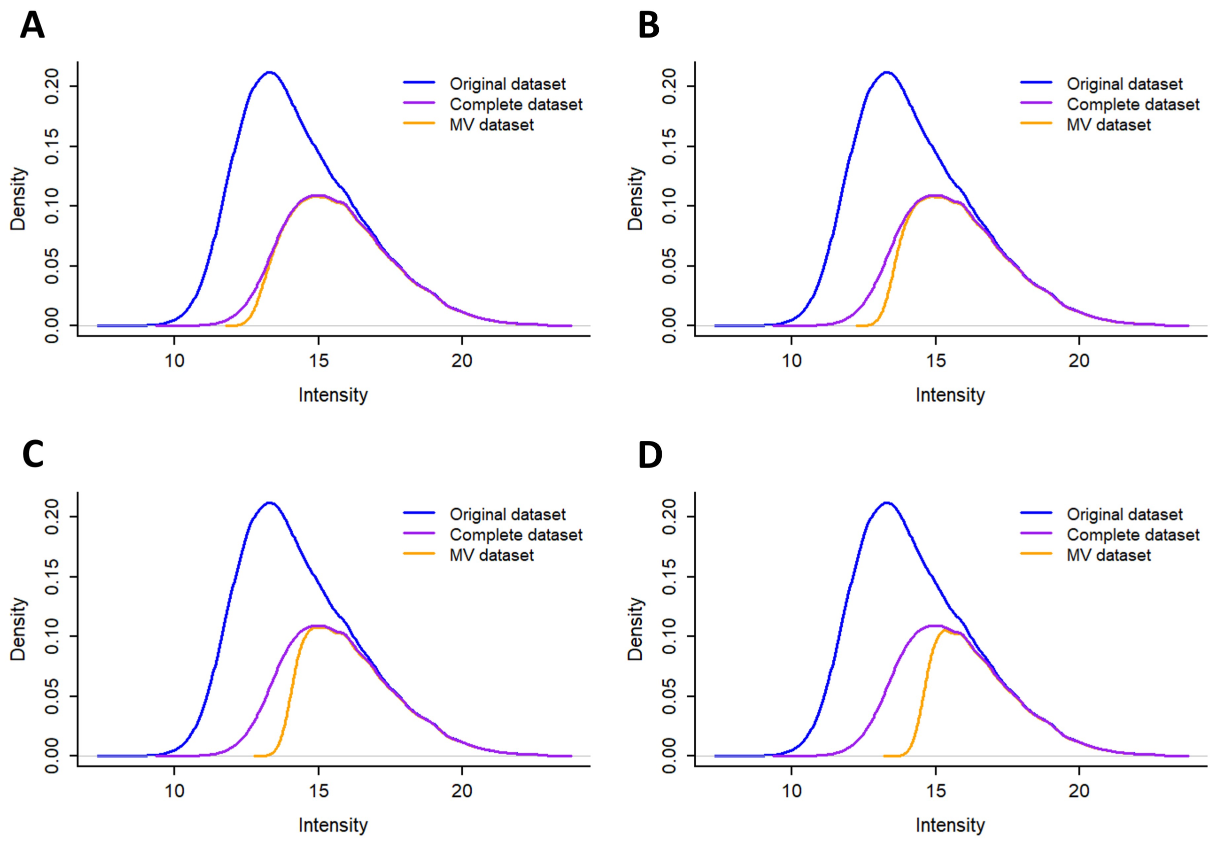


**Figure S3.** Data distribution under cutoffs and MCAR at missingness levels of (a) 5%, (b) 10%, (c) 20%, and (d) 30%. The blue curve shows the original distribution with missing values; the purple curve shows the distribution after removing proteins with missing values; and the yellow curve shows the distribution after introducing cutoffs and MCAR missingness into the complete dataset.

#### Figure S4: Precision–Recall curves under cutoff and MCAR missingness at varying proportions.

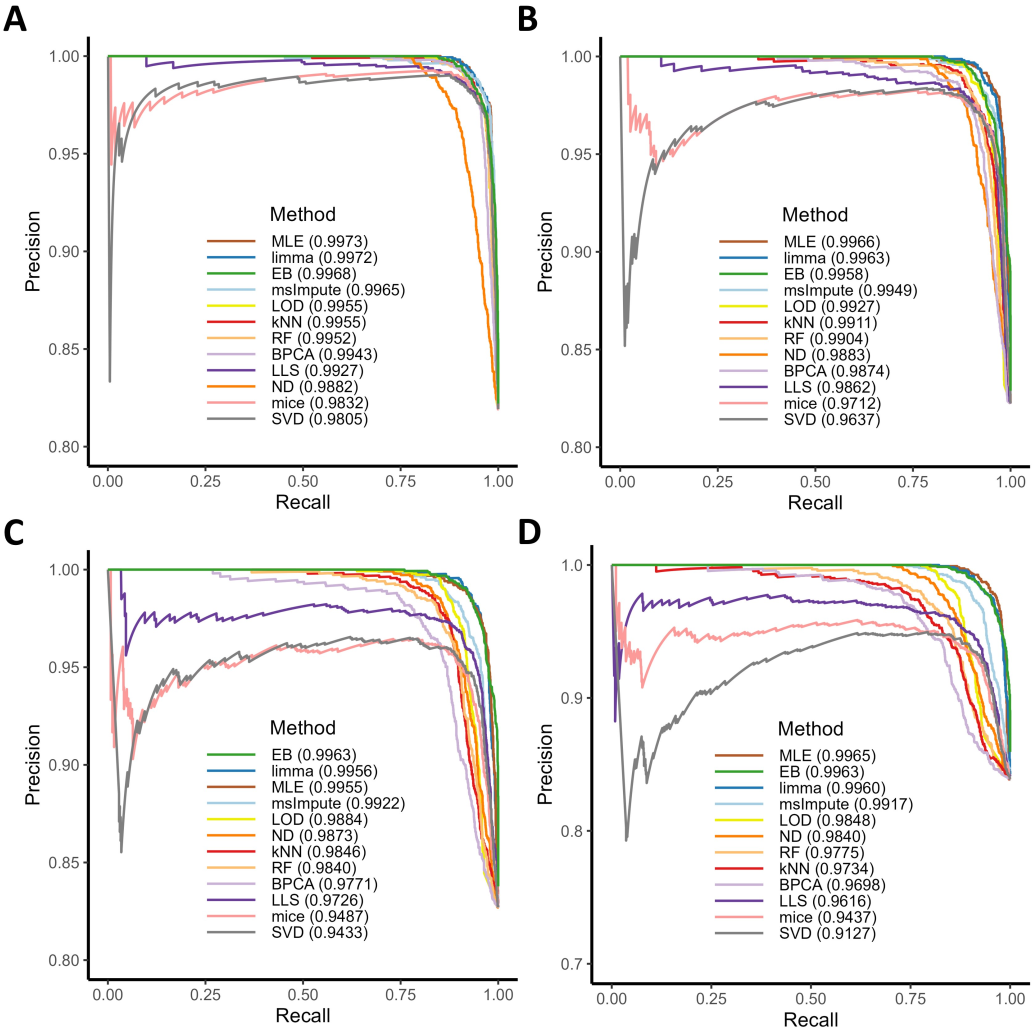


**Figure S4.** Precision–Recall curves under cutoffs and MCAR missingness at varying missingness levels. (A) 5% missingness. (B) 10% missingness. (C) 20% missingness. (D) 30% missingness. The area under the precision–recall curve (AUPRC) is reported in the legend. True positives are defined as proteins with $p < 0.05$ under the standard $t$-test, and false positives as proteins with $p \geq 0.05$.

#### Figure S5: Predictive fit of dataset with 20% values set to missing under cutoffs and MCAR missingness.

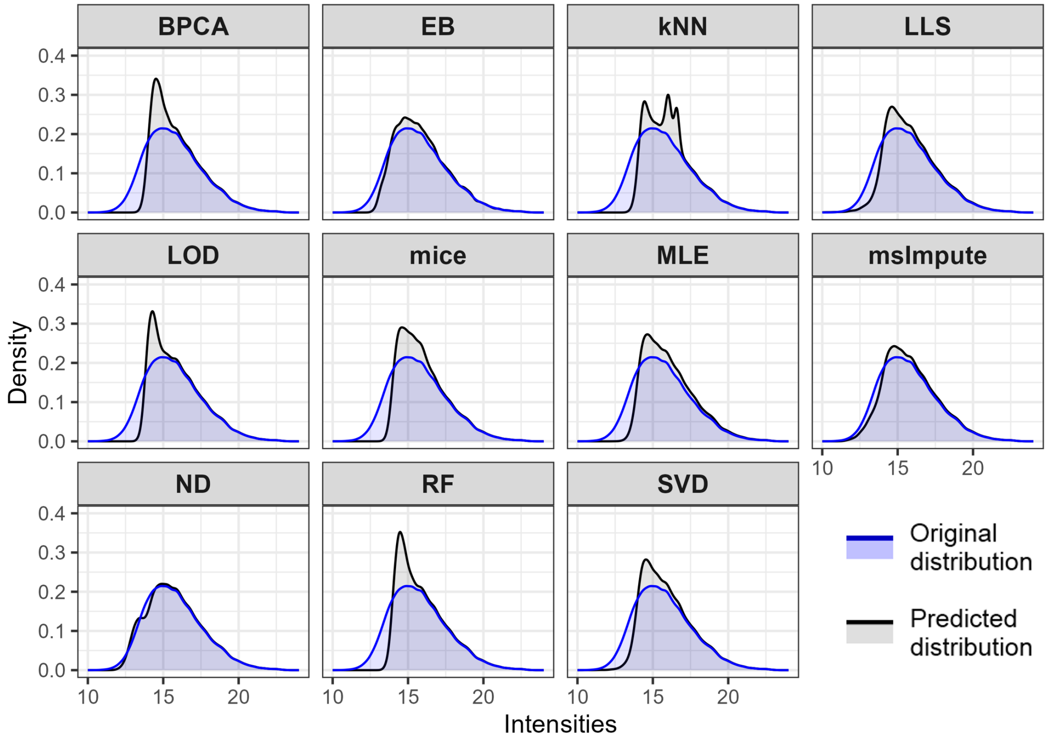


**Figure S5.** Predictive fit for a dataset with 20% values artificially set to missing under cutoffs and MCAR mechanism. The blue curve shows the distribution of the original (complete) data, and the black curves show the distributions predicted by each method. For EB, predicted values were generated from the posterior predictive distribution.

### Figure S6: Example of MCMC diagnostic plots

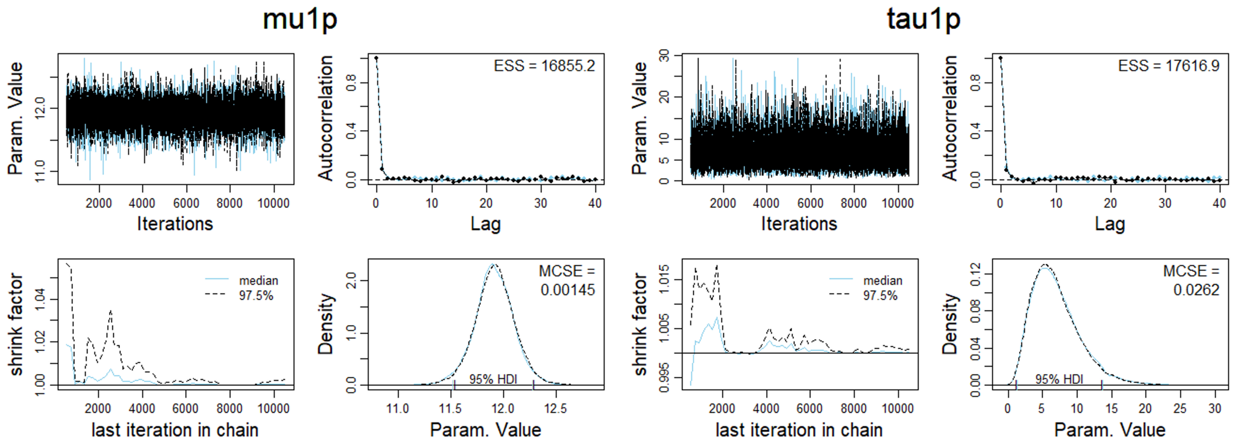


**Figure S6.** MCMC diagnostic plots for the group 1 mean and group 1 precision of protein ZN316_HUMAN in the CQE-A dataset. The upper-left panel shows the trace plots for two chains, which overlap closely, indicating good mixing. The upper-right panel shows the autocorrelation function (ACF), where correlations drop rapidly toward zero for lags greater than one, suggesting minimal autocorrelation. The lower-left panel displays the ratio of between-chain to within-chain variance; values close to 1 indicate good convergence (analogous to $\hat{R}$). The lower-right panel shows the posterior density of the parameter, demonstrating consistent posterior distributions across chains. Diagnostic figures were generated using code adapted from John K. Kruschke's Doing Bayesian Data Analysis (2nd edition).
